# Massively Parallel Analysis of Human 3′ UTRs Reveals that AU-Rich Element Length and Registration Predict mRNA Destabilization

**DOI:** 10.1101/2020.02.12.945063

**Authors:** David A. Siegel, Olivier Le Tonqueze, Anne Biton, Noah Zaitlen, David J. Erle

## Abstract

AU-rich elements (AREs) are 3′ UTR cis-regulatory elements that regulate the stability of mRNAs. Consensus ARE motifs have been determined, but little is known about how differences in 3′ UTR sequences that conform to these motifs affect their function. Here we use functional annotation of sequences from 3′ UTRs (fast-UTR), a massively parallel reporter assay (MPRA), to investigate the effects of 41,288 3′ UTR sequence fragments from 4,653 transcripts on gene expression and mRNA stability. The library included 9,142 AREs, and incorporated a set of fragments bearing mutations in each ARE. Our analyses demonstrate that the length of an ARE and its registration (the first and last nucleotides of the repeating ARE motif) have significant effects on gene expression and stability. Based on this finding, we propose improved ARE classification and concomitant methods to categorize and predict the effect of AREs on gene expression and stability. Our new approach explains 64±13% of the contribution of AREs to the stability of human 3′ UTRs in Jurkat cells and predicts ARE activity in an unrelated cell type. Finally, to investigate the advantages of our general experimental design for annotating 3′ UTR elements we examine other motifs including constitutive decay elements (CDEs), where we show that the length of the CDE stem-loop has a significant impact on steady-state expression and mRNA stability. We conclude that fast-UTR, in conjunction with our analytical approach, can produce improved yet simple sequence-based rules for predicting the activity of human 3′ UTRs containing functional motifs.

## 1 Introduction

3′ untranslated regions play an important role in regulating mRNA fate by complexing with RNA binding proteins that help control mRNA localization, translation, and stability [1, 2, 3]. Identification of a consensus UUAUUUAU sequence in the 3′ UTRs of human and mouse mRNAs encoding tumor necrosis factor (TNF-*α*) and a variety of other inflammatory mediators led to the suggestion that these AU-rich elements AREs) could be important for regulating gene expression [4]. Subsequent studies confirmed that these and other AREs interact with ARE-binding proteins such as AUF1 (also known as hnRNPD), HuR and other Hu family proteins, and the CCCH zinc finger-containing RBPs ZFP36 (tristetraprolin), ZFP36L1, and ZFP36L2 [5], to alter mRNA degradation and protein expression [6]. In most cases, AREs have been reported to destabilize mRNAs, although in some cellular contexts certain AREs and ARE-binding proteins have been shown to stabilize mRNAs [6, 7]. Subsequent analyses of the human genome concluded that as many as 58% of human genes code for mRNAs that contain AREs [8, 9, 10], suggesting that these elements play a major role in regulating expression of a large group of genes.

An initial classification of AREs was proposed based upon studies of human 3′ UTR sequences and analyses of mutation effects and the activity of synthetic AREs [11]. AUUUA motifs were recognized as critical for destabilizing effects of many 3′ UTRs, and in many cases destabilizing AREs contained two or more overlapping repeats of this motif. However, some AUUUA motif-containing sequences were not destabilizing and some AU-rich sequences that lacked the AUUUA motif had potent mRNA destabilizing activity. Based upon these observations, Chen and Shyu [11] divided AREs into two classes of AUUUA-containing AREs and a third class of non-AUUUA AREs. Class I AUUUA-containing AREs had 1-3 copies of scattered AUUUA motifs coupled with a nearby U-rich region or U stretch, whereas class II AUUUA-containing AREs had at least two overlapping copies of the nonamer UUAUUUA(U/A)(U/A) in a U-rich region. Non-AUUUA AREs had a U-rich region and other unknown features, and the relationship of these sequences to AUUUA-containing AREs remains poorly understood. Subsequent studies based on analyses of a set of 4884 AUUUA-containing AREs led to a new classification based primarily on the number of overlapping AUUUA-repeats [8, 9, 10]. This classification system, with five clusters distinguished by the number of repeats, was used to identify AUUUA-containing AREs in the human genome. AREs identified using this classification were found to be abundant in 3′ UTRs of human genes. However, the functional activities of this large set of ARE-containing 3′ UTR sequences remains mostly unknown.

To address this shortcoming we relied on novel experimental and analytical strategies. First, we leveraged Massively Parallel Reporter Assays (MPRAs), which are capable of simultaneously characterizing the regulatory impact of thousands of mRNA sequence fragments for functional analysis [12, 13, 14, 15, 16]. We previously developed a massively parallel method for functional annotation of sequences from 3′ UTRs (fast-UTR) [15]. In that study we used fast-UTR to analyze a set of 3000 160 nt sequences from human 3′ UTRs that were highly conserved across mammalian species and confirmed that AREs and constitutive decay elements (CDEs) in these sequences are important contributors to 3′ UTR-mediated mRNA destabilization. However, due to the limited number of sequences included in that study, we were unable to identify rules that governed the activity of sequences that conformed to these motifs.

To address this issue, in this work we designed, produced, and analyzed a fast-UTR library containing a comprehensive set of ARE sequences from human mRNA 3′ UTRs in human cell lines. We focused largely on Jurkat T cells, given the importance of AREs in T cell development [5] and autoimmunity. We also studied BEAS2B airway epithelial cells to begin to investigate whether key findings were mimicked in a different cell type. In total, the MPRA in our current study included 41,288 sequences from 4,653 transcripts. Applying novel analytic approaches to this set of sequences [17] allowed us to identify rules that predict functional effects of AREs on mRNA stability and stead-state expression.

To verify that our combined fast-UTR approach and analytic strategy was not uniquely successfully in the context of AREs, we also considered a second important class of 3′ UTR regulatory elements known as constitutive decay elements (CDEs) [4, 18]. CDEs are conserved stem loop motifs that bind to the proteins Roquin and Roquin2, resulting in increased mRNA decay [18]. CDEs include an upper stem-loop sequence of the form UUCYRYGAA flanked by lower stem sequences. Lower stem sequences are formed by 2-5 nt pairs of reverse-complementary sequences (e.g. CCUUCYRYGAAGG has a lower stem length of 2). We similarly generated a fast-UTR library containing a comprehensive set of CDEs. We again obtained novel sequence based rules predicting CDE function. Finally, to show the potential of our approach to aid functional interpretation of arbitrary 3′ UTR sequence fragments, we conclude by developing statistical models to predict the activity of entire 160 nt 3′ UTR sequence fragments in our full MPRA data set.

## 2 Materials and Methods

### 2.1 Sequence Design

#### 2.1.1 Definition of 3′ UTR regions and regulatory regions

We segmented 3′ UTRs from human RefSeq transcripts (v68) into 160 nt sliding windows with a shift of 80 nt. Only regions of at least 20 nucleotides were included; segments shorter than 160 nt were padded with a sequence from the CXCL7 3′ UTR (NM_002704, 475-602) that had minimal regulatory effects in prior experiments [15]. We then identified those 160 nt 3′ sequence segments that contain suspected ARE motifs and CDE motifs, as well as several other features that are described in the Supplementary Methods. The ARE motifs targeted for inclusion were those defined according to the ARED website [8] as of Fall 2014, generally containing one or more repeat of AUUUA, following rules that are precisely defined in the Supplementary Methods (Table S1). Since the design of our experiment, the ARED group has revised their ARE class definitions into “clusters” in a more recent publication [10], and our analysis follows those updated conventions. The CDE motifs contain a central sequence of UUCYRYGAA, surrounded by a “lower stem” of 2-5 bp and then an unpaired nt.

In addition to the reference regions containing ARE and CDE motifs, we designed segments that contain mutated versions of these motifs. For AREs, we mutated the central U of the AUUUA pentamer to a C or G, alternating down the sequence (for example AUUUAUUUA was mutated to AUcUAUgUa). Since several features were targeted for inclusion, we noticed that there are several regions in the dataset that contain core AUUUA pentamers that were shorter than the ARED definitions (for instance, a region containing a CDE might also contain an “AUUUA” within the 160 nt window), so we have designed segments where 429 of these smaller AUUUAs were mutated into AUcUAs [8]. For CDEs, two types of mutations were introduced: 1) UUCYRYGAA was mutated to UagYRYGAA, and 2) the CDE sequence was shuffled. In total, the assay contained 41,288 segments from 13,334 3’UTR regions from 4,653 RefSeq transcripts.

### 2.2 Fast-UTR assay

An oligonucleotide pool containing the full set of library segments was produced by massively parallel synthesis (Agilent), amplified by PCR, and cloned into the BTV lentiviral plasmid as previously described [15]. Barcodes were introduced using random 8-mer sequences incorporated into the PCR primer in order to obtain systematic estimates of technical variability. A lentiviral library was used to transduce Jurkat T cells or BEAS2B airway epithelial cells expressing tetracycline transactivator tTA, allowing for doxycycline-regulated reporter transcription. Cells were maintained in culture for 2 weeks after transduction. Cells were left untreated (t_0_) or treated with doxycycline (1 *μ*g/ml) for 4 hours to inhibit reporter transgene expression (t_4_) prior to isolation of DNA and RNA using the AllPrep DNA/RNA/Protein Mini Kit (Qiagen# 80004) according to the manufacturers protocol. RNA was reversed transcribed to cDNA, and both genomic DNA and cDNA were amplified to produce sequencing libraries. Sequencing was performed using an Illumina HiSeq 4000. An average of at least 20 clones per segment were obtained for each time point for each cell line to obtain systematic estimates of the technical variability.

### 2.3 Data Analysis

#### 2.3.1 Steady State Expression and Stability

We quantify mRNA activity in two ways, by measuring “steady-state expression” and “mRNA stability”. To quantify steady state expression we use the count-normalized ratio before the addition of doxycycline:

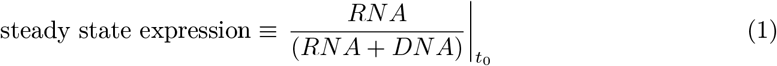

To quantify mRNA stability, we use a ratio of ratios, comparing the expression 4 hours after the addition of doxycycline (t_4_) to the steady-state expression (t_0_):

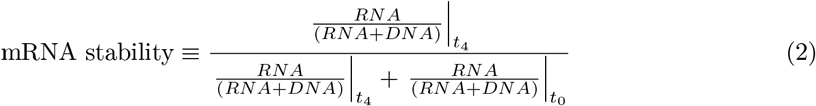

For example, if a segment contains a binding site for an mRNA degrading protein, the steady state expression and stability will be small; whereas a segment containing a binding site for a stabilizing protein will result in a large estimate of steady state expression and stability.

To investigate the effect of a mutation on the activity of a UTR segment we examine the change (Δ) in steady state expression or stability by subtracting the reference value from the mutant value: Δ ≡ Mutant − Reference. If a functional element (ARE or CDE) is destabilizing, then mutating it will likely lead to a positive change in stability and expression by disrupting the corresponding mRNA binding proteins.

#### 2.3.2 GC Content

GC content has been suggested to impact measurements of gene expression and stability [19, 20]. Because the primary purpose of this study is to examine the relative contributions of known sequence motifs to the expression and stability of mRNAs, confounding effects of GC content are a critical consideration.

Therefore, to account for GC-content in the fast-UTR data, we fit 5th-order polynomials to the steady state expression and stability as a function of GC-content, and subtract them from the steady state expression and stability. We assessed the performance of *N*th order polynomial fits (*N* ∈ {1, …, 7}) through leave-one-out cross validation and found that the prediction accuracy (Pearson correlation between predicted and actual steady state expression and stability) did not improve significantly after *N* = 5. See Supplement for further details.

### 2.4 Evaluation of ARE Classification Methods

To examine the quality of previously proposed as well as novel classification systems for AREs, we propose assessment via prediction quality. We assess the quality of predictions by determining correlations between measured and predicted RNA expression and stability values using a leave-one-chromosome-out approach. Broadly, we argue that better classification approaches will more accurately predict the affect of individuals AREs.

Leaving out one chromosome eliminates some sources of bias, such as generating predictions from overlapping genomic regions. Therefore we split the data into a training set of 3′ UTR segments from 23 chromosomes with which we generated our model parameters, and a test set of segments from the 24th chromosome (treating Y as a separate chromosome from X). We repeated this process for each chromosome to generate a prediction for every ARE-containing segment in the dataset. When predicting the effect of a designed mutation on a mutant and reference sequence pair, we only consider mutations that disrupt pentamers of the ARE. For categorical predictions, we calculated the categorical means from the training set, then used those means as predictors for the test set. For example, if the mean for category 3 AREs was 0.4 in the training set, we assigned a “predicted value” of 0.4 to any category 3 ARE in the test set. For regression-based methods we performed the regressions on the training set to generate model parameters, then applied those models to the test set to produce predictions. The correlation between predicted and measured values is then reported.

The rules for four ARE prediction methods are detailed below, and additional methods are given in the Supplement. Unless stated otherwise, a segment is classified by its longest ARE if more than one is present:

i. ARE Plus: We used the five cluster motifs described by Bakheet, Hitti, and Khabar [10] to create training and test categories. Cluster 1 and 2 motifs total 13 nucleotides, with AU-rich segments flanking one or two AUUUA core motifs, respectively. Clusters 3, 4 and 5 include 3, 4, or 5 exact AUUUA repeats respectively. This system differs somewhat from the earlier ARED definitions described by the same group [8] which we used for the initial design.
ii. Naive Effective Length Pentamers: Pentamers classified by the “effective length” according to the formula floor((length(nt) + registration − 2)/4). “Registration” refers to the starting nucleotides of the ARE within the initial AUUUA pentamer: an ARE that starts AUUU*=0, UUUA*=1, UUAU*=2, and UAUU*=3. No mismatches allowed.
iii. Effective Length (nt): AREs classified by the “effective length” (length(nt) + starting registration), rather than class of pentamer. No mismatch allowed.
iv. Lasso K-Mer Regression: Each 160 nt segment was broken down into a list of 156 5-mers, which were used in a Lasso [21] regression scheme following Reference [14], solving the linear model y=Xb+c for effect sizes b (y is the outcome vector containing steady state expression or stability, and X is the frequency matrix). Further details are given in the Supplement. Here the training set is not limited to segments with AREs, and includes every segment in the full MPRA; but the test set is limited to the same set of segments with AREs as Figure 3 (any AUUUA pentamer with length 6 or higher, plus the length-5 pentamer “AUUUA”). To predict the effect of mutations, we simply subtract the predicted expression or stability of the wild-type from the predicted expression or stability of the mutant; we do not train and test on the difference data directly.

## 3 Results

### 3.1 An MPRA to Investigate the Effect of AREs on Gene Expression and Stability

We used our previously developed fast-UTR MPRA to test effects of a large set of human 3′ UTR segments containing elements conforming to previously defined ARE motifs on mRNA steady-state levels and mRNA stability (Figure 1). In our initial analysis, we observed a significant effect of GC content of the 3′ UTR segments on steady state expression, with increased GC content associated with reduced expression (Figure S1A). We recently reported a similar finding in a fast-UTR-based analysis of ≈27,000 70-nt RNA binding protein binding 3′ UTR sequences tested in mouse primary T cells [20]. GC content also had an apparent effect on mRNA stability (Figure S1B). The GC content effects are not due to AU-rich elements, since their effect is to do the opposite: AU-rich elements have low GC-content but generally lower steady-state expression and stability. In subsequent analyses, we adjusted for GC content as described in the Materials and Methods (Table S2 and Figure S1).

**Figure 1:**
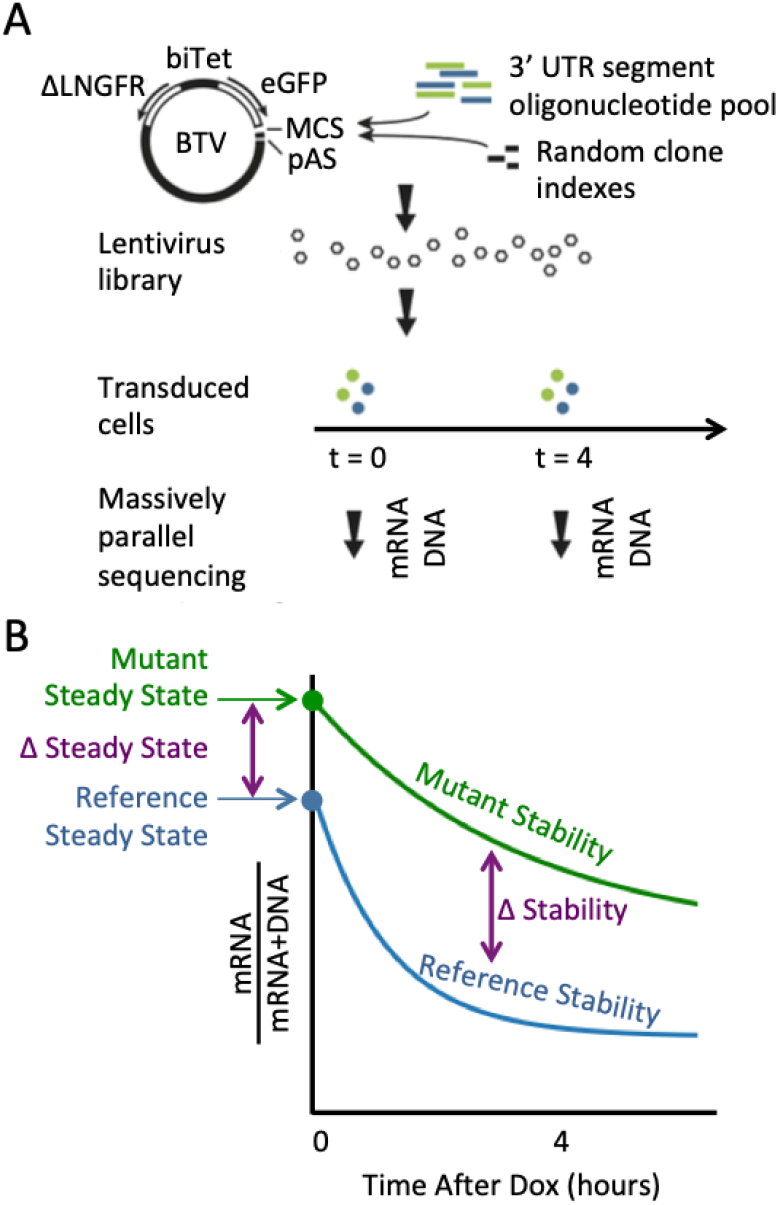
The fast-UTR MPRA. (A) The BTV plasmid includes a bidirectional tetracycline regulated promoter (biTet) that drives expression of enhanced green fluorescent protein (EGFP) and a reference protein (truncated low-affinity nerve growth factor receptor, ΔLNGFR). The EGFP reporter transgene includes a multiple cloning site (MCS) for insertion of 3′ UTR test sequences and a polyadenylation signal (pAS). Pools of 160-mer oligonucleotides containing 3′ UTR segments were inserted into BTV together with random octamer indexes used to identify each clone. Cells were transduced with BTV lentiviral libraries and massively parallel sequencing was used to measure 3′ UTR segment sequences in genomic DNA and mRNA isolated from cells. (B) Steady state mRNA levels were determined from clone read counts for mRNA samples before the addition of doxycycline (Dox). mRNA stability was estimated from mRNA read counts obtained before and 4 h after the addition of Dox to inhibit transcription. The blue line represents a 3′ UTR segment with an element that promotes rapid mRNA decay and the green line represents a sequence with an inactivating mutation of the destabilizing element that increases steady-state mRNA levels and reduces the decay rate.

Measurements were performed at several time points with respect to the addition of doxycy-cline: before dox was added (*t*_0_), and 4 hours after dox. 4 hours was chosen because evaluating RNA/(RNA+DNA) at *t*_4_/(*t*_4_ + *t*_0_) showed greater sequence-based variation than using *t*_2_ or *t*_6_ in preliminary data. To set a minimum threshold for the data quality of a 3′ UTR sequence segment, we required segments to be represented by at least 5 clones with more than 5 counts of DNA each, and for each segment to have at least 1 count of RNA in at least 1 clone. While low RNA counts are generally indicative of low gene expression, we believe the measurement of zero RNA counts to most likely be a technical artifact, since the correlations across sequencing replicates are greatly increased when segments with zero RNA counts are removed.

### 3.2 Established Categories of ARE Predict Gene Expression and Stability

We found that ARED-Plus classification was associated with functional activity of 3′ UTR segments. When 3′ UTR segments were classified according to the presence of sequences conforming to the ARED clusters, ARE cluster membership had a significant impact on both steady state RNA level (Figure 2A) and RNA stability (Figure 2B). Both steady state level and stability decreased as the number of ARE repeats increased. Segments that contained the minimal AUUUA sequence only (without any ARED-Plus motifs) had a slightly lower steady state level and stability than segments with no AUUUA or ARED-Plus motif (a two-sided Welch’s t-test give *p* = 7 × 10^−6^, 4 × 10^−30^, 7 × 10^−3^ and 3 × 10^−23^ for Figures 2A-D respectively). There was a consistent decrease in steady state RNA and stability with cluster number from clusters 1 through 3. Cluster 5 motif-containing segments were more active than cluster 3; cluster 4 segments were also associated with low expression and stability although the number of segments in this cluster was small and the confidence interval was larger than for other clusters.

**Figure 2:**
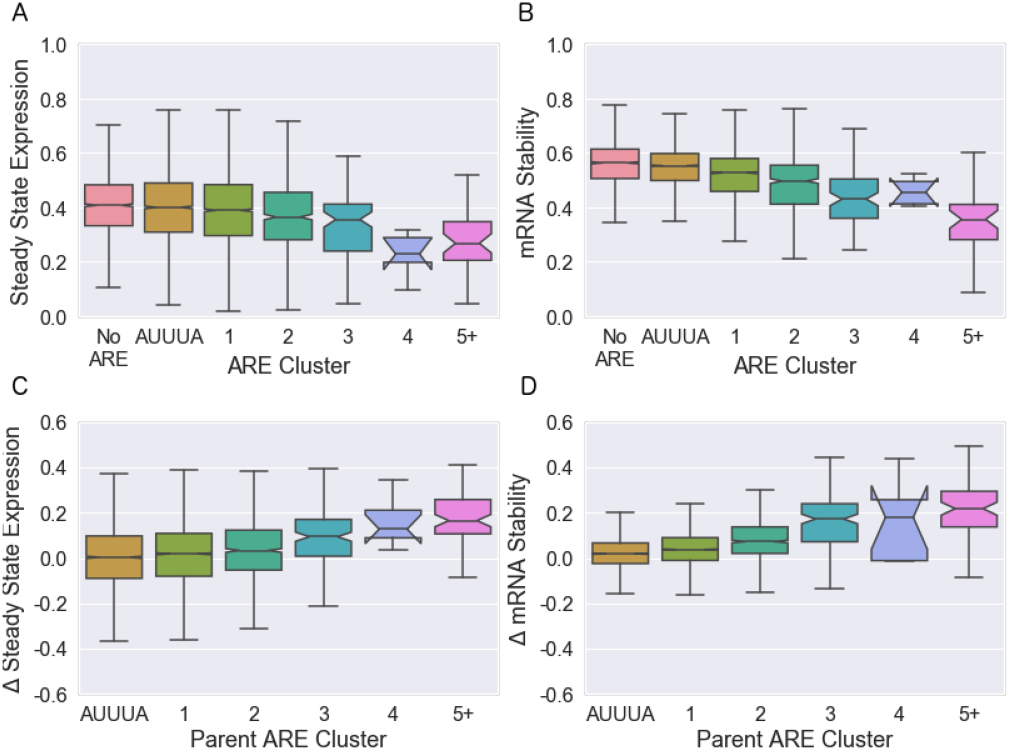
Effect of established ARE clusters on mRNA expression and stability in Jurkat T cells. Boxplots show medians and quantiles, and notches indicate 95% confidence intervals. (A, B) Effects of 3′ UTR segments containing ARED motifs on (A) steady-state RNA expression and (B) RNA stability. (C, D) Effects of mutations in ARED motifs on (C) steady-state expression and (D) RNA stability. Δ represents values for segments containing mutations in the ARE motif minus values for the corresponding reference segment with intact ARE motifs. For the 3% of segments with more than one ARE, we categorize them according to the largest ARE present. The “AUUUA” cluster consists of segments that contain the minimal “AUUUA” sequence but lack the flanking sequences required for ARED-Plus. ARE cluster membership was significantly associated with changes in all four measures (*p* = 3 × 10^−22^ for A, 6 × 10^−149^ for B, 1 × 10^−31^ for C, and 8 × 10^−177^ for D by linear regression).

The 160-nt 3′ UTR regions that we tested contained both AREs and surrounding sequences that could also affect reporter expression. To more directly examine the effects of the AREs on expression levels and RNA stability, we leveraged our novel technology to systematically examine the effect of mutations that disrupted the AREs (Figure 2C and D). As expected, disrupting AREs increased steady state RNA and stability. The effects of mutations were clearly related to the number of AUUUA repeats, as represented by ARED-Plus cluster.

Although there was a clear relationship between ARED-Plus cluster and the effects of mutating AREs, there was considerable variation that was not explained by ARED-Plus cluster. We suspected that this might be due to limitations of existing cluster definitions, but it might also be attributable to technical variability inherent in the assay.

To address this issue, we developed MPRAudit, a novel method to determine the fraction of variance explained by sequence variation in MPRAs and other barcoded assays [17]. The premise of MPRAudit is that the technical variation from sequence to sequence can be estimated from the variation from clone to clone for a given sequence segment. If this technical variability can be determined, then we attribute the remaining variability between sequences to be due to sequence variation. When MPRAudit is applied to pairs of segments where the ARE motif has been deliberately mutated, it can determine the fraction of variance caused by the type of mutation and the flanking sequence. We call this quantity the “explainability”, denoted *b*^2^. When *b*^2^ is close to 0, technical variance is the source of all variation. When *b*^2^ is close to 1, the variation from sequence to sequence is the primary source of variation and technical sources are small. Table 1 shows that there is a substantial amount of sequence variation remaining within each of these established groups. This suggests that there are biologically different categorical subgroupings within each currently defined cluster of AREs that can explain more of the effects of ARE-containing sequences than established clusters.

**Table 1:** Fraction of variance explained by sequence variation within the clusters of AREs defined using ARED-Plus motifs [10]. If *b*^2^ is zero then the variation is due to technical factors; if *b*^2^ is greater than zero then the data support more refined subgroupings of AREs with different functional properties within each class.

| ARED-Plus Cluster | Fraction of Variance Driven by Sequence Variation |  |  |  |
| --- | --- | --- | --- | --- |
| | $b^2 \Delta$ Steady State | $b^2 \Delta$ Stability | N<sub>Seqs</sub> Steady State | N<sub>Seqs</sub> Stability |
| AUUUA | $0.2269 \pm 0.0006$ | $0.057 \pm 0.002$ | 4098 | 3714 |
| Cluster 1 | $0.2271 \pm 0.0004$ | $0.116 \pm 0.001$ | 5278 | 4725 |
| Cluster 2 | $0.214 \pm 0.002$ | $0.307 \pm 0.005$ | 1122 | 1025 |
| Cluster 3 | $0.32 \pm 0.01$ | $0.41 \pm 0.02$ | 158 | 155 |
| Cluster 4 | $0.09 \pm 0.15$ | $0.16 \pm 0.21$ | 9 | 9 |
| Cluster 5 | $0.35 \pm 0.02$ | $0.50 \pm 0.01$ | 77 | 73 |

### 3.3 ARE Registration and Length Affect mRNA Stability

To create a new ARE classification system that could explain more of the observed effects of ARE-containing sequence segments, we examined three parameters that we hypothesized would correlate with ARE activity: the length of the ARE, the starting registration of the ARE, and the conservation status of the ARE. For this analysis we looked only at perfect matches to a repeating “AUUUA” pattern of length at least 6, plus exact matches to the “AUUUA” pentamer itself. Hence we examined “UUAUUU”, despite the fact that it does not contain an “AUUUA”, but did not include “UAUUU” because it is too short.

As expected, the length of the ARE was associated with activity. We found a gradual increase in ARE activity as a function of the ARE length (in nucleotides) without obvious abrupt increases at 5-nt (AUUUA pentamer) intervals (Figures 3A and B show mRNA stability; Figures S2A and S2B show steady state). This suggests that the length of the ARE may provide additional information beyond that provided by the ARED-Plus clusters, which depend on the number of pentamers.

**Figure 3:**
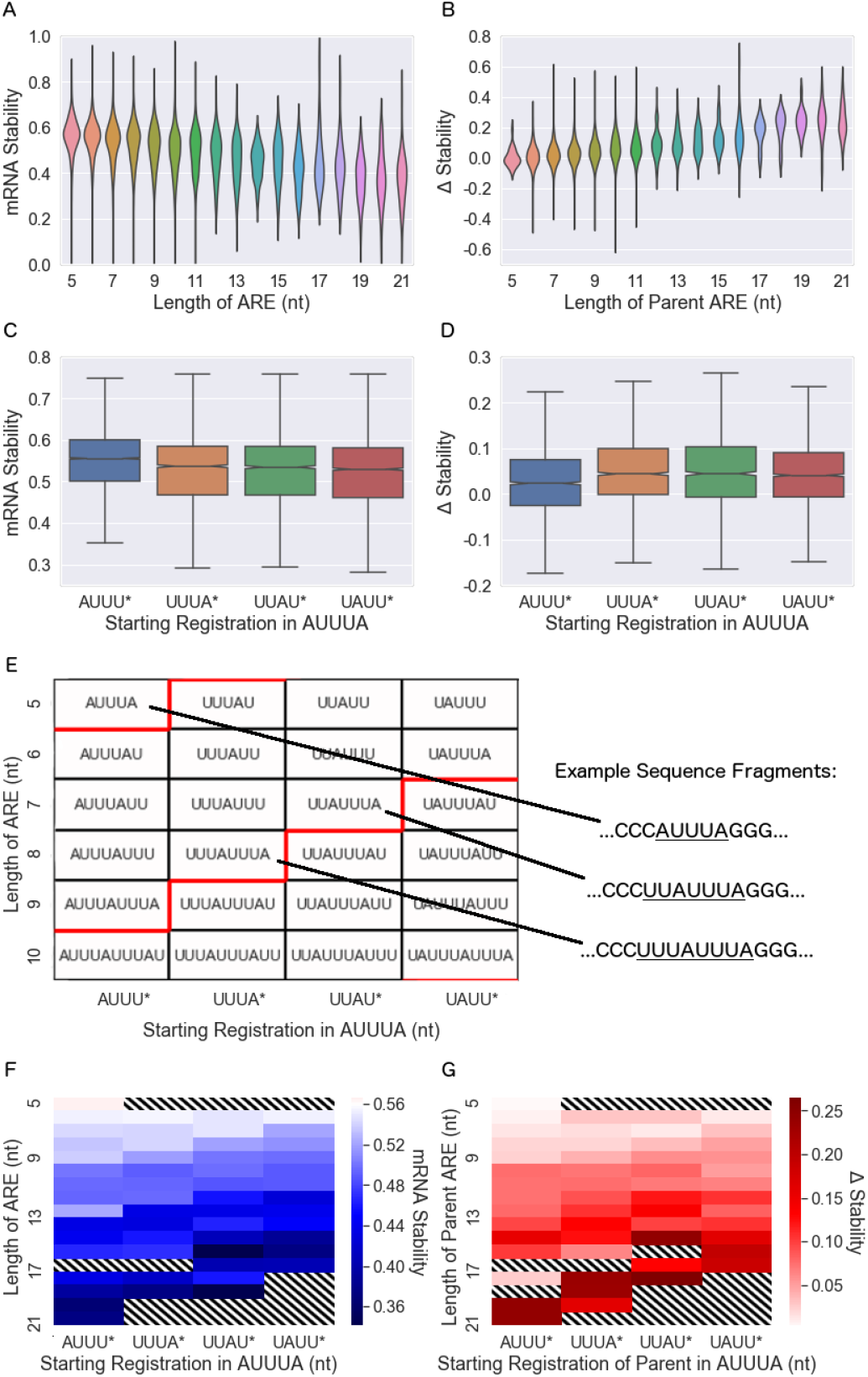
mRNA stability is associated with ARE length and registration. (A, B) Length of the ARE (in nt) is associated with mRNA stability for the segment containing the ARE (A) and with the change in stability that resulted from mutation of the ARE (B). (C.D) Starting registration of the ARE is associated with mRNA stability (C) and mutation-induced change in stability (D). Notches in the boxplots that show confidence intervals are barely visible at this scale. Linear regression shows that the slopes of these datasets (treating the x-axis as a continuous variable) are nonzero, with two-sided p-values of 5 × 10^−302^, 5 × 10^−269^, 1 ×10^−44^, and 1 ×10^−5^, respectively. (E) Classification of AREs by length and registration. The x-axis gives the starting registration of the ARE, while red lines indicate the step between ending registrations of *UUUA (boxes above the red line) and *UUAU (boxes below the red line). (F) mRNA stability according to length and registration. For comparison, the mean mRNA stability for segments lacking an ARE is 0.56 (represented as white). (G) Mutation-induced change in mRNA stability according to length and registration. Standard errors for each entry in each panel are given in the Supplement (Figs S4 and S5). In panels F and G, boxes that weren’t studied, boxes that had no corresponding segments in the dataset, and boxes with large standard errors (≥0.07) are excluded and filled with diagonal stripes.

We next considered the starting point of the ARE within the AUUUA pentamer, which we term the “starting registration”. We define an ARE starting with A to have starting registration 0, and each time a nucleotide is removed from the front the registration increases by 1. For instance, the sequence “AUUUAUU” has length 7 and registration 0, while “UUUAUUU” has length 7 and registration 1 (and we consider this an ARE despite the absence of an “AUUUA”). Since overlapping pentamers have a periodicity of four nucleotides, an ARE may have one of four possible starting registrations (and one of four possible ending registrations, see Figures S6 and S7). We found that starting registration did have an effect on ARE activity, with sequences beginning with “AUUU” being less active than sequences with other registrations (Figures 3C and D show mRNA stability; Figures S2C and S2D show steady state). We conclude that both ARE length and registration have significant associations with ARE activity in univariate analyses.

In contrast to ARE length and registration, we found that conservation status had little effect on ARE activity. We used phastCons, which identifies conserved elements from 100 vertebrate species [22], to classify AREs as conserved (overlap with phastCons conserved regions) or non-conserved. Surprisingly, there was little difference between the activity of conserved and non-conserved AREs (Fig. S5). We also considered whether more highly conserved sequences might be more active than other conserved sequences. However, within the set of conserved sequences, we found no significant association of the phastCons lod score with ARE activity in our fast-UTR assay.

Since ARE length and registration were each independently associated with ARE activity, we explored the relationship between these two parameters further. Any sequence of AUUUA pentamer repeats can be classified by its length in nucleotides and its starting (or ending) registration. Figure 3E shows the sequence motifs of several example AREs, organized according to ARE length along the y-axis and ARE starting registration along the x-axis. Motifs of constant ending registration appear along the diagonals; for instance, the boxes directly above (or below) the red lines have the same ending registration. Figures 3F and 3G are similar to 3E, except the color of each box represents the mean stability across sequence segments or change in stability with mutation, respectively. As expected, activity increases with increasing ARE length (moving from top to bottom of heat maps). The diagonal contours arise due to effects of registration. Similar effects are apparent when analyzing steady state RNA levels rather than stability, and when using ending registration rather than the starting registration as a parameter (Supplementary Results and Figure S6 and S7). These results suggest that registration, in additional to length, should be an integral part of ARE classification and provide insights into underlying mechanisms.

## 4 Better Classification Methods for AREs

We anticipated that our enhanced understanding of how the length and registration of AREs affect gene expression and stability would allow us to make improvements to existing categories of AREs and to create better prediction methods. To quantify the performance of these categories, we used leave-one-chromosome-out cross validation to train and test predictions for each of the segments with AREs in our dataset. Leaving out one chromosome avoids overfitting from overlapping 3′ UTR segments, since overlapping segments will be within the same chromosome.

We examined correlations between out-of-chromosome predictions and measured data for several classifications of AREs and linear models as described in the Methods section. Using the 5 clusters defined by ARED-Plus [10] (as in Figure 2A and B) led to a modest correlation with steady state mRNA level and a somewhat higher correlation with mRNA stability (Table 2). As expected, correlations were somewhat higher for mutation-induced changes in mRNA level and stability, since these measures depend more critically on the targeted ARE itself than on the other sequences within the 3′ UTR segments. Including both ARE length (in nt) and registration to form an “effective length” (as suggested by Figure 3E-G) resulted in significant increases in correlations, again with the exception of mutation-induced changes in steady-state level. To verify statistical significance, we compared the squared out-of-chromosome residuals generated by the ARED-Plus categories to the squared out-of-chromosome residuals generated by the “effective length” method. A Mann-Whitney U test finds that the out-of-chromosome residuals were smaller for the “effective length” categories with p-values of *p* = 0.013, 1.6 × 10^−21^, and 8.6 × 10^−9^ for steady state mRNA, stability, and change in stability due to mutation, respectively. There was no apparent difference for change in steady state mRNA due to mutation (p = 0.5). We also considered a set of related methods based on ARE length, starting or ending registration, and allowance of a mismatch to the strict “AUUUA” motif, but found none that outperformed the “effective length” (length and starting registration) method (Table S5).

**Table 2:** Out-of-chromosome correlations between predictions and measured data for existing and novel ARE categories (i-iii) and a prediction method (iv). See Table S5 for further categories and S6 for Beas2B data.

| Out-Of-Chromosome Correlation Between Predictions and Measured Data |  |  |  |  |  |
| --- | --- | --- | --- | --- | --- |
| # | Category or Method | Steady-State Expression | Stability | $\Delta$ Steady-State | $\Delta$ Stability |
| i | ARED-Plus Clusters | 0.09 | 0.21 | 0.14 | 0.29 |
| ii | Naive Effective Length Pentamers | 0.11 | 0.29 | 0.14 | 0.34 |
| iii | Effective Length (nt) (Length + Registration) | 0.14 | 0.30 | 0.14 | 0.36 |
| iv | Lasso K-Mer Regression (K=5) | 0.26 | 0.49 | 0.13 | 0.34 |

The finding that the activity of an ARE is highest when it starts and ends with UAUU* and *UUAU, respectively, is consistent with studies [23, 24, 25] that claim “UAUUUAU” to be the “minimal” ARE motif, since UAUUUAU is the smallest ARE that starts and ends with both UAUU* and *UUAU. However, we observe that the effects of some shorter AREs, while smaller in magnitude, still remain statistically significant. In fact, Figure 3 shows that even AREs with no “AUUUA” (the motifs UUUAUU, UUAUUU, and UUUAUUU) can have a statistically significant effect on gene stability: compiling data from segments with these three motifs, we find that their stability is 0.013±0.002 lower than the experimental mean with no ARE. The effect of mutations (Figure 3G) are also significant for each of these motifs: the change in stability is 0.030±0.009, 0.030±0.011, and 0.019±0.009 for UUUAUU, UUAUUU, and UUUAUUU, respectively. This suggests that “AUUUA” may not be absolutely required for ARE binding and function.

The wealth of data generated by this protocol also opens the possibility of more complicated sequence-based prediction approaches to 3′ UTR function. As a first step in this direction, we report the results of a method that makes use of k-mer decomposition and regression. This use of k-mers is different from the other methods we have attempted, in that it is agnostic to the presence of AREs in the training data and might be sensitive to the presence of other active elements in the 3′ UTR aside from AREs. This additional information led the k-mer method to have the best performance on predictions of steady state expression and stability, of all the methods we tried. However, its performance on predicting the change in steady state and stability with ARE mutation was slightly worse than the performance of the effective length method, perhaps because it was not specifically targeting AREs. The distribution of k-mer effect sizes, the identity of top k-mers, and the effect sizes of ARE-related k-mers are given in Figures S3 and S4. AUUUA is one of the most negative 5-mers, and the other ARE component sequences (UUUAU, UUAUU, UAUUU) have negative lasso amplitudes as well. On the other hand, substituting a C or G for one of the nucleotides of these four ARE component sequences results in fewer negative amplitudes: only 2 out of 40 of the mutated 5-mers are negative (*p* = 1 × 10^−4^ using a Fisher exact test). The mutated 5-mers are also more positive than the 980 remaining 5-mers in the dataset (*p* = 0.003 using a chi-square test with Yates’ correction for continuity).

Overall, these results show that the new knowledge introduced by our analysis is helpful for categorizing and classifying AREs and does a better job of predicting gene expression and the effects of mutations than previously established categories.

Table 2 shows that our methods improve predictions, but Table 3 shows that there is still room for further improvement. MPRAudit allows us to calculate an upper limit to the performance of prediction methods, and also to calculate the fraction of variance explained by ARE categories, out of the total possible variance in the dataset caused by sequence variation (removing the fraction caused by known technical factors). It does this by calculating the fraction of sequence variation that remains within each group (restricting the analysis to sequences with AREs) before and after the groupings of Table 2 are applied. We do not apply this technique to the method of k-mers because the method of k-mers does not create groupings and it has a very large number of variables. We find that our predictions of steady state expression and stability explain a small fraction of the total variation caused by differences in sequence, which might be expected by the relatively small correlations in the first three columns of Table 2 (the squared correlation is related to the fraction of variance explained). On the other hand, our novel groupings explain up to two thirds of the sequence-based variation for the effects of ARE mutations on sequence stability, as the technical sources of variation make up a sizeable fraction of the total variance. All told, we conclude that human 3′ UTRs have many additional unknown regulatory mechanisms, and ARE-mediated decay is just one contributer to 3′ UTR effects on mRNA stability.

**Table 3:** Fraction of sequence-driven variance explained by given classification systems, as calculated by MPRAudit. *b*^2^ = 1 would imply a perfect model of ARE behavior. The *b*^2^ statistic is first calculated for the entire dataset, ignoring groupings, then calculated within groups, and compared. The fraction explained by categories is calculated as 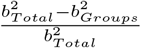.

| Categorization | $b^2$ Expression | $b^2$ Stability | $b^2 \Delta$ Expression | $b^2 \Delta$ Stability |
| --- | --- | --- | --- | --- |
| Bakheet Pub 2 | $0.00 \pm 0.02$ | $0.05 \pm 0.02$ | $0.05 \pm 0.09$ | $0.39 \pm 0.12$ |
| Effective Length Pentamers | $0.02 \pm 0.02$ | $0.06 \pm 0.02$ | $0.06 \pm 0.09$ | $0.54 \pm 0.13$ |
| Effective Length | $0.03 \pm 0.02$ | $0.07 \pm 0.02$ | $0.08 \pm 0.09$ | $0.64 \pm 0.13$ |

## 5 CDE Steady-State Expression and Stability Vary with Stem Length

To determine if the fast-UTR approach could also be applied to other known 3′ UTR elements we next considered CDEs. The fast-UTR library we constructed contained all 345 3′ UTR segments with sequences conforming to a previously-defined degenerate CDE stem-loop motif [18]. We found that the destabilizing effects of the CDE motif increased with increasing stem length (Figure 4).

**Figure 4:**
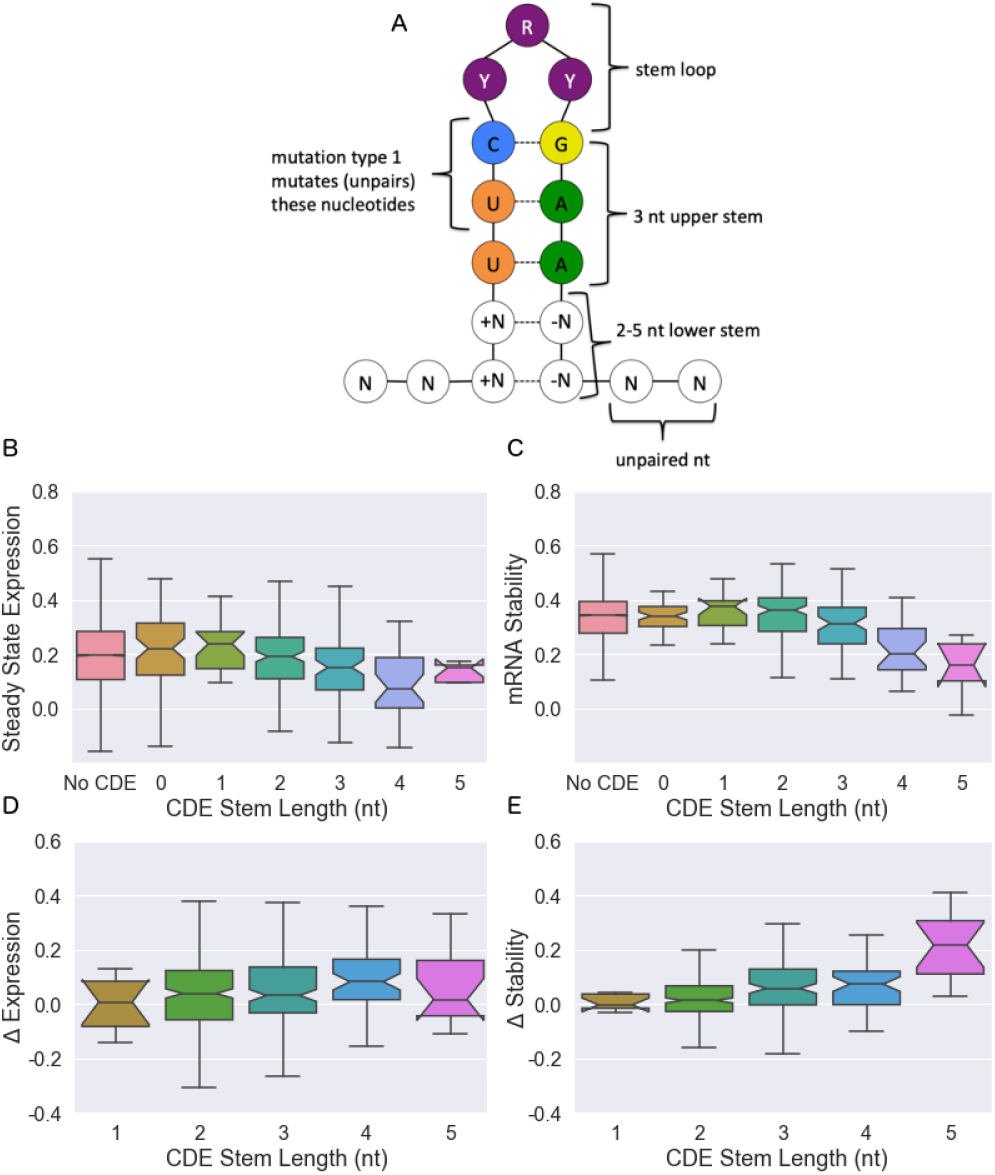
Effect of CDE stem length on mRNA expression and stability. (A) Schematic of the CDE stem loop in our investigation. (B) Steady-state expression, (C) stability, (D) change of steady-state expression with CDE mutation, and (E) change of stability with CDE mutation. Boxplots give medians and quantiles, notches denote 95% confidence intervals. While lower stem lengths of 2-5 nt are considered active, CDE motifs with stem lengths of 0-1 are also contained within the dataset and shown here for reference. Linear regression gives slopes that differ from zero with *p* = 1.2 × 10^−6^, 2.8 × 10^−7^, 3.2 × 10^−3^, and 2.9 × 10^−13^, for panels (b-e), respectively.

As with AREs, we deliberately introduced mutations into the CDE sequences to isolate the effects of these motifs from the overall sequence. For CDEs, two types of mutations were introduced: 1) the central motif UUCYRYGAA was mutated to UagYRYGAA, or 2) the entire motif and surrounding stem were shuffled. For both cases, the differences between mutant and reference sequences were recorded. Since these two types of mutations had similar effects on gene expression and stability (Figure S9), we combined them for further analysis. We found that the effect of CDE mutation increased with the length of the outer stem (Figures 4C and D), consistent with the effects on steady state and stability measurements in Figures 4A and B: increasing the length of the CDE stem leads to a decrease in steady-state expression and the stability of the mRNA, and the effects of mutations increase with increasing stem length.

## 6 Other Regulatory Elements

Although our analysis focuses on the behavior of AREs and CDEs, Table 3 suggests that there may be many other regulatory elements in the human 3’ UTR. To investigate additional possibilities, we analyzed the motifs that were highlighted by Rabani et al [14] as being stabilizing or destabilizing in zebrafish and that also happened to be present in our dataset. Figure S10A and B show that only a few of these motifs are active in human cell lines, most notably the Pumilio and miR430 motifs (in addition to AREs). miR430 does not exist in humans, but the human miRNA miR-302a shares the same seed sequence [26] and therefore might account for this finding. Comparison with Figures S10C and D show that GC-residualization plays an important role in determining the activity of these sequences.

## 7 AREs Are More Active in Beas2B Cells Than Jurkat

To determine whether insights obtained from our studies of Jurkat T cells would apply to another cell type, we used fast-UTR to study the same 3′ UTR segment library in Beas2B human bronchial epithelial cells. Application of the full set of methods tested for Jurkat T cells showed that the “effective length” model was also the best model for explaining ARE activity in Beas2B cells (Figure S7 and Table S5). However, Figure 5 shows that the magnitude of ARE effects differed between cell types, with Beas2B cells showing a larger response than Jurkat cells to the presence of AREs in a 3′ UTR segment. As a function of the length of the ARE, gene expression decreases more rapidly, decay decreases more rapidly, and the effects of mutations are larger for Beas2B cells. The Beas2B standard errors are larger than the Jurkat standard errors because fewer of the Beas2B sequences passed our quality control filters. As a result, the ARE predictions have higher correlations with experimental data for the Beas2B data than the Jurkat overall, but the improvement of our effective length categorization over previous categorizations is smaller, due to the decrease in quality of the training data.

**Figure 5:**
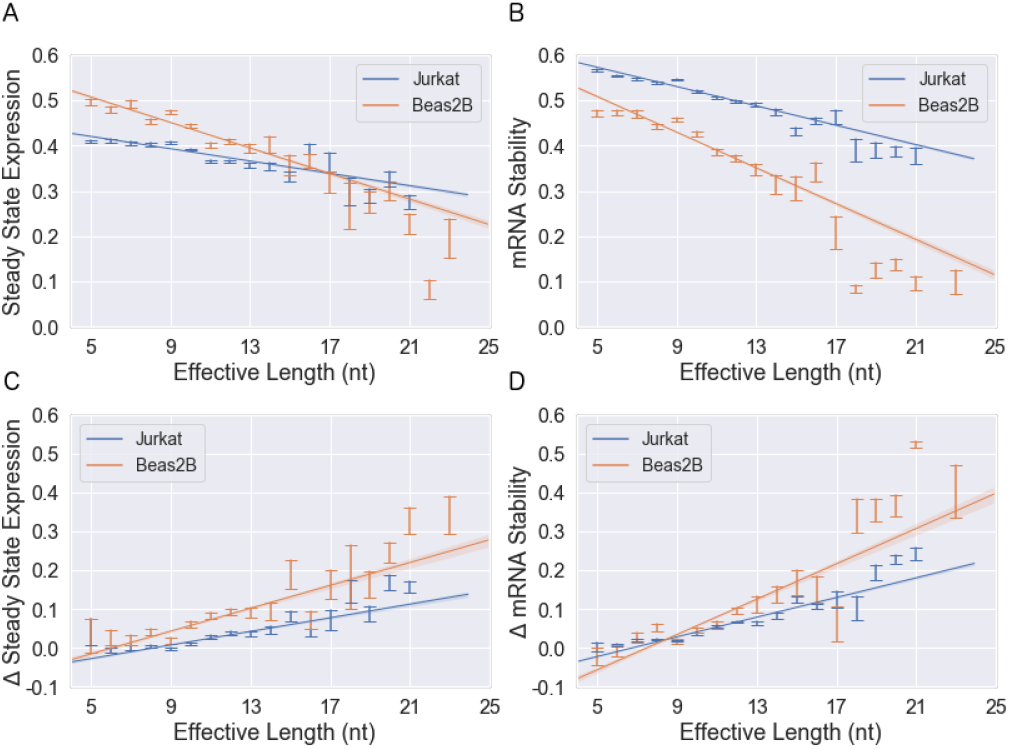
Comparison of ARE activity between Jurkat and Beas2B. (A) Steady state expression, (B) stability, (C) change in steady state expression due to deliberate mutations, and (D) change in stability due to deliberate mutations, as a function of the ARE “effective length” (ARE length plus starting registration). Error bars give standard errors. Linear fits are shown to each dataset as a guide to the eye. AREs are more active in Beas2B in each case, with 2-sided p-values of 9 × 10^−21^, 3 × 10^−34^, 7 × 10^−70^, and 3 × 10^−14^, respectively [27].

## 8 Discussion

In this work we have developed and analyzed a massive experimental system for examining 3′ UTR biology. Our approach uncovered novel features that affect the stability and steady-state expression of AREs and CDEs in 3′ UTRs. We show that the length of an ARE, as well as the starting or ending nucleotide, has an effect on gene expression and stability, and verify these findings through designed mutations of the active motif. In CDEs we similarly show that longer stem loops have an effect on the activity of the motif. Using our recently developed method MPRAudit, we show that a model consisting of ARE length and registration explains up to 64% of the effects of mutations on the stability of AREs.

There are several conclusions that the results of Tables 2 and S5 allow us to draw: adding the length and registration of the ARE to form an “effective length” improves the prediction correlation by almost 50% for steady state expression and stability; but the effect of mutations on steady state expression remains difficult to predict. We found that allowing one mismatch generally gives worse performance than requiring a perfect ARE match. Lasso regression on k-mers performs best on predictions of expression and stability, since it incorporates features from outside the ARE; but it makes slightly worse predictions for the effects of targeted ARE mutations than our methods that rely on ARE length and registration alone.

The steady state expression and stability of CDEs are consistent with our understanding of their secondary structure. If the central motif of the CDE (UUCYRYGAA) is a binding site for destabilizing proteins, increasing the prominence of its stem loop could have an impact on the binding affinity of the CDE.

This research has many other limitations that suggest directions for future work. We analyze a subset of 3′ UTRs and not the whole genome; and we focus on the behavior of regulatory sequences, but ignore the important contribution of secondary structure. Although we study the behavior of two cell lines, we have not investigated primary cells or performed in vivo studies. Within this dataset, a significant amount of variance is explained by the categories of sequences that we have uncovered, but a significant amount of variance remains unexplained by our work. While our work has produced some general rules for predicting the steady state expression and stability of mRNAs, we note that there are many ARE binding proteins and we are unable to disentangle their effects from one another. By making the raw data in this work publicly available, we hope machine learning researchers, statisticians, and geneticists will make further improvements to the models we have devised here.

## 9 Acknowledgements

DAS, AB and NZ were funded by NIH grants K25 HL121295, U01 HG009080, R01 HG006399, R01 CA227237, R03 DE025665, R01 CA227466, R01 ES029929, R01 GM110251, R01 HL124285, and DoD grant W81XWH-16-2-0018

DJE and OLT were funded by NIH grants R35 HL145235, R01 GM110251, and R01 HL124285.

The authors would like to thank Wenxue Zhao, Mark Ansel, Steven Brenner, and Ilias Soares for useful discussions.

## References

[1] Mayr, C. Regulation by 3′-untranslated regions. Annual Review of Genetics 51, 171–194 (2017).

[2] Castello, A. et al. Insights into RNA biology from an atlas of mammalian mRNA-binding proteins. Cell 149, 1393–1406 (2012).

[3] Glisovic, T., Bachorik, J. L., Yong, J. & Dreyfuss, G. RNA-binding proteins and post-transcriptional gene regulation. FEBS Letters 582, 1977–1986 (2008).

[4] Caput, D. et al. Identification of a common nucleotide sequence in the 3-untranslated region of mRNA molecules specifying inflammatory mediators. Proceedings of the National Academy of Sciences 83, 1670–1674 (1986).

[5] Hodson, D. J. et al. Deletion of the RNA-binding proteins ZFP36l1 and ZFP36l2 leads to perturbed thymic development and t lymphoblastic leukemia. Nature Immunology 11, 717–724 (2010).

[6] Barreau, C. AU-rich elements and associated factors: are there unifying principles? Nucleic Acids Research 33, 7138–7150 (2005).

[7] Peng, S. S.-Y. RNA stabilization by the AU-rich element binding protein, HuR, an ELAV protein. The EMBO Journal 17, 3461–3470 (1998).

[8] Bakheet, T. ARED: human AU-rich element-containing mRNA database reveals an unexpectedly diverse functional repertoire of encoded proteins. Nucleic Acids Research 29, 246–254 (2001).

[9] Bakheet, T. ARED 2.0: an update of AU-rich element mRNA database. Nucleic Acids Research 31, 421–423 (2003).

[10] Bakheet, T., Hitti, E. & Khabar, K. S. A. ARED-plus: an updated and expanded database of AU-rich element-containing mRNAs and pre-mRNAs. Nucleic Acids Research 46, D218–D220 (2017).

[11] Chen, C.-Y. A. & Shyu, A.-B. AU-rich elements: characterization and importance in mRNA degradation. Trends in Biochemical Sciences 20, 465–470 (1995).

[12] Sample, P. J. et al. Human 5′ UTR design and variant effect prediction from a massively parallel translation assay. Nature Biotechnology 37, 803–809 (2019).

[13] Avsec, Ž. et al. Deep learning at base-resolution reveals motif syntax of the cis-regulatory code (2019).

[14] Rabani, M., Pieper, L., Chew, G.-L. & Schier, A. F. A massively parallel reporter assay of 3′ UTR sequences identifies in vivo rules for mRNA degradation. Molecular Cell 68, 1083–1094.e5 (2017).

[15] Zhao, W. et al. Massively parallel functional annotation of 3′ untranslated regions. Nature Biotechnology 32, 387–391 (2014).

[16] Kreimer, A. et al. Predicting gene expression in massively parallel reporter assays: A comparative study. Human Mutation 38, 1240–1250 (2017).

[17] Siegel, D. A., Tonqueze, O. L., Biton, A., Erle, D. J. & Zaitlen, N. Mpraudit quantifies the fraction of variance describedby unknown features in massively parallel reporter assays. bioRxiv (2020).

[18] Leppek, K. et al. Roquin promotes constitutive mRNA decay via a conserved class of stem-loop recognition motifs. Cell 153, 869–881 (2013).

[19] Benjamini, Y. & Speed, T. P. Summarizing and correcting the GC content bias in high-throughput sequencing. Nucleic Acids Research 40, e72–e72 (2012).

[20] Litterman, A. J. et al. A massively parallel 3′ UTR reporter assay reveals relationships between nucleotide content, sequence conservation, and mRNA destabilization. Genome Research 29, 896–906 (2019).

[21] Tibshirani, R. Regression shrinkage and selection via the lasso. Journal of the Royal Statistical Society. Series B (Methodological) 58, 267–288 (1996).

[22] Siepel, A. Evolutionarily conserved elements in vertebrate, insect, worm, and yeast genomes. Genome Research 15, 1034–1050 (2005).

[23] Wiklund, L., Sokolowski, M., Carlsson, A., Rush, M. & Schwartz, S. Inhibition of translation by UAUUUAU and UAUUUUUAU motifs of the AU-rich RNA instability element in the HPV-1 late 31 untranslated region. Journal of Biological Chemistry 277, 40462–40471 (2002).

[24] Zubiaga, A. M., Belasco, J. G. & Greenberg, M. E. The nonamer UUAUUUAUU is the key AU-rich sequence motif that mediates mRNA degradation. Molecular and Cellular Biology 15, 2219–2230 (1995).

[25] Lagnado, C. A., Brown, C. Y. & Goodall, G. J. AUUUA is not sufficient to promote poly(a) shortening and degradation of an mRNA: the functional sequence within AU-rich elements may be UUAUUUA(u/a)(u/a). Molecular and Cellular Biology 14, 7984–7995 (1994).

[26] Rosa, A., Spagnoli, F. M. & Brivanlou, A. H. The miR-430/427/302 family controls mesendo-dermal fate specification via species-specific target selection. Developmental Cell 16, 517–527 (2009).

[27] Paternoster, R., Brame, R., Mazerolle, P. & Piquero, A. Using the correct statistical test for the equality of regression coefficients. Criminology 36, 859–866 (1998).

