## Supplement for "Massively Parallel Analysis of Human 3′ UTRs Reveals that AU-Rich Element Length and Registration Predict mRNA Destabilization" (from another version of this preprint: version 2, 2020-02-18)

#### Supplementary Methods

##### Library preparation and sequencing

###### Definition of 3′ UTR segments

Segments were defined using 160 nt sliding windows with a shift of 80 nt. Only segments of at least 20 nucleotides were included; sequences shorter than 160 nt were padded with as many nt of a standard padding sequence as required to bring the test sequence length to 160 nt. The standard padding sequence was derived from a segment of the 3′ UTR of CXCL7 previously shown to have minimal activity in our assay [1]: UUUGUUCUG UUUCUGCCAAACUUCUUUAACUCCCAGGAAGGGUAGAAUUUUGAAACCUUGAUUUUCUAGAGUU CUCAUUUAUUCAGGAUACCUAUUCUUACUGUAUUAAAAUUUGGAUAUGUGUUUCAUUCUGUCUC AAAAAUCACAUUUUAUUCUGAGA.

###### Selection of segments containing predicted regulatory elements

A total of 13,334 3′ UTR segments of 4,653 RefSeq genes as defined in the UCSC database (v68) were selected based upon meeting one or more of the following criteria:

- AREs: AREs were identified based upon the classification proposed for the ARE database (see Table 1)
- CDEs: UUCYRYGAA surrounded by a "lower stem" of 2-5 bp and then an unpaired nt: 345 segments

A set of other segments that did not contain the ARE or CDE motifs listed above were also included and were used for comparison. These included 3′ UTR segments with single nucleotide polymorphisms associated with altered gene expression (eQTLs)[2] and 3′ UTR segments from genes that have been associated with asthma in GWAS[3]. In addition to 3′ UTRs designed based on the RefSeq reference sequences, we included designed mutations (lower case indicates mutated nucleotides) and naturally occurring variants as follows:

- ARE: AUUUAUUUA -> AUcUAUgUa (mutate the central U of the first AUUUA pentamer to a c, the next central U to a g, etc.)
- Minimal "AUUUA" sequences (not conforming to the ARE database motifs): AUUUA -> AUcUA
- CDE: 1) UUCYRYGAA -> UagYRYGAA and 2) random shuffles of the motif and stem
- Pumilio: 1) UGUAAAUA -> UuugAAUA, 2) UGUACAG -> UuUgCAG, and 3) random mutations

*These authors contributed equally to this work

- TargetScan predicted binding sites for selected miRNAs (let-7/98, miR-15/16/195/424/497, miR-17-5p/20/93.mr/106/519.d, miR-27ab, miR-29abc, miR- 25/32/92/92ab/363/367,miR-26ab/1297): random mutations in the first 6 nucleotides of the miRNA motif.

Note that all 3′ UTR segments containing AREs and CDEs were mutated, whereas for minimal "AUUUA", Pumilio and miRNA targets, only those segments that were selected based on the selection criteria listed above were included (reference and mutant versions).

Naturally occurring variants:

- We included all naturally occurring variants within the selected 3' UTR segments that were observed in the 1000 Genomes pilot 1 study (1,092 individuals) [2] or the Consortium on Asthma among African-ancestry Populations in the Americas (CAAPA) [3]. A total of 6733 of the 3′ UTR segments had one or more variants.

Many segments contain multiple regulatory elements, and not every segment had significant numbers of counts after sequencing.

During the time between library preparation and analysis a new set of ARE class definitions were published by Bakheet et al[4], given in Table 1. We used these updated definitions for our data analysis.

Notably, the prior system used lower numbers to refer to AREs with more repeats, whereas the updated system uses higher numbers to refer to AREs with more repeats (e.g. AREs with 5 repeats were categorized as Group I instead of V).

#### Plasmids

pSIN4-EF2-ABCG2-IRES-Neo was a gift from Ren-he Xu (Addgene plasmid # 25983, Watertown, MA) [1]. pMA2818 was a gift from Mikhail Alexeyev, PhD of the University of South Alabama College of Medicine[5]. The pSIN4_tTA_advanced plasmid was created by cloning the EcoRI-BamHI fragment containing the tTA advanced sequence into the EcoRI-BamHI digested pSIN4_ABG2-IRES-Neo. The plasmid was then sequence verified by Sanger Sequencing.

The BTV plasmid was previously described [6]. pMDLg/pRRE was a gift from Didier Trono (Addgene plasmid # 12251) [7]. pMD2.G was a gift from Didier Trono (Addgene plasmid # 12259). pRSV-Rev was a gift from Didier Trono (Addgene plasmid # 12253).

#### Production of fast-UTR library

We used oligonucleotide pools that were produced by massively parallel synthesis (Agilent). 41443 Oligonucleotides (0.2 pmol) were amplified by two rounds of PCR. The first PCR (Q5 polymerase ; 98°C for 45s followed by 6 cycles of 98°C for 15s, 60°C for 30s and 72°C for 30s followed by 72°C for 60s) was done using the Oligo_FW and Oligo_RV primers (see Primer Table). 1 $\mu$L of the first PCR was used as template for the second PCR (Q5 polymerase; 98°C for 45s followed by 7 cycles of 98°C for 15s, 60°C for 30s and 72°C for 30s followed by 72°C for 60s) using the Subset_FW and Subset_RV primers. PCR products were purified using a PCR purification kit (Qiagen, #28104, Germantown, MD) and digested with PacI (NEB #R0547L, Ipswich, MA) and MluI-HF (NEB #R3198L) in CutSmart buffer for 2h at 37°C. The digested products were purified from a 1% Tris-acetate-EDTA gel with a Zymoclean Gel DNA Recovery Kit (Zymo Research # D4007, Irvine, CA) and ligated into BTV plasmid with Quick DNA ligase (NEB # M2200S). Ligation mixtures were purified with the QIAquick PCR Purification Kit (Qiagen #28104) and introduced into E. cloni (Lucigen # 60052-2, Middleton, WI) by electroporation to produce a mean of at least 30 clones per oligonucleotide ($1.2 \times 10^6$ clones in total). After overnight culture at 37°C in LB supplemented with ampicillin, plasmids were extracted using a Maxiprep kit (Zymo Research # D4203).

#### Lentivirus production

Lentiviruses were produced by co-transfecting Lenti-X 293T cells with the BTV reporter plasmid pMDLg/RRE, pMD2g and pRSV-rev (Addgene# 12253) at a molar ratio of 3:1:1:1 with TransIT-293 (Mirus # MIR 2705,

Madison, WI) according to the manufacturer's protocol. After 24h the medium was discarded and replaced with fresh medium for another 48h. Medium was collected and concentrated with Lenti-X concentrator (Takara # 631232, Mountainview, CA). Lenti-X concentrator was added to conditioned medium at the ratio of 1:3 v:v and was then rotated at 4°C for 1h to overnight. The viruses were spun at 1500g for 45min at 4°C and the pellet was then resuspended in RPMI culture medium (100x). If mentioned otherwise the virus were produced by the UCSF Viracore. Briefly, HEK293T cells were seeded at 70,000 cells per $cm^2$ in 15 cm tissue culture dishes in 20 mL media (DMEM, 10% FBS) and incubated overnight at 37 °C, 5% $CO_2$. 24 h after plating, 12 $\mu$g of lentiviral transfer vector was transfected together with 7 $\mu$g psPAX2 (Addgene #12260) and 3 $\mu$g pMD2.G (Addgene #12259) with 50 $\mu$L jetPRIME transfection reagent (Polyplus) according to the manufacturer's protocol. 72 h post-transfection, lentiviral supernatant was collected and passed through 0.45 $\mu$m filters (Millipore). Aliquots were stored at −80 °C.

#### Lentivirus titering

The functional titer of purified lentiviral particles was determined by transduction of Jurkat E6 cells with serial dilutions of lentivirus preparations FACs analysis of the LV reporter 72hrs post-transduction as described in [8].

#### Cell culture

Jurkat T cells were grown in RPMI (Life Technologies #A1049101, Carlsbad, CA) supplemented with 10% Fetal Bovine Serum (Life Technologies #10437-028) with 1x penicillin/streptomycin (Sigma-Aldrich #P4333, St. Louis, MO). BEAS2B cells were grown in DMEM/F12 (Life Technologies #11320082) supplemented with 10mM HEPES (Hyclone #SH30237.01), 2mM L-Glutamine (Life Technologies #25030-081), 1X Non-Essential Amino Acid (GenClone #25-536), 1x penicillin/streptomycin (Sigma-Aldrich #P4333), 10% Bovine Calf Serum (Hyclone #SH3007304, Logan, UT). Lenti-X 293T cells (Clontech # 632180, Mountain View, CA) were grown in DMEM (Life Technologies #11995065), supplemented with 10mM HEPES (Hyclone #SH30237.01), 2mM L-Glutamine (Life Technologies #25030-081), 1x penicillin/streptomycin (Sigma-Aldrich #P4333), Bovine Serum (Life Technologies #10437-028). Cell lines were authenticated by STR profiling at the University of Arizona genetics core and treated with 25$\mu$g/mL plasmocin (InvivoGen #ant-mpt, San Diego, CA) to eliminate Mycoplasma contamination.

#### Clonal isolation of tTa cell lines

The tTA-IRES-neo lentivirus were the used to transduced Jurkat and Beas2B cells. 24h after transduction JurkatG418 at, respectively, 500$\mu$g/mL and 250$\mu$g/mL. Single clones were then screened for tTA activity by limiting dilution method.

#### Transduction and purification

Lentiviral transduction was performed at a multiplicity-of-infection of approximately 0.3. After 72h incubation with lentivirus, transduced cells were enriched using the MACSelect LNGFR microbeads kit (Miltenyi #130-091-330). Cells were washed and magnetically labeled using MACSelect LNGFR MicroBeads and loaded onto an LS Column (Miltenyi #130-042-40, Bergisch Gladbach, Germany) placed in a MidiMACS Separator. After washing, magnetically labeled transfected cells were recovered from the column and analyzed by flow cytometry using the APC Labeling Check reagent (Miltenyi #130-098-892). Cells were maintained for 2 weeks prior to use in fast-UTR assays.

#### Fast-UTR assays

$120 \times 10^6$ cells per time point were harvested 0h and 4h after addition of doxycycline (1$\mu$g/mL) to the medium and genomic DNA (gDNA) and RNA were isolated using the AllPrep DNA/RNA/Protein Mini Kit (Qiagen# 80004) according to the manufacturer's protocol.

#### Sequencing libraries

We prepared both genomic DNA and RNA sequencing libraries for all samples. For genomic DNA, we used the genomic DNA isolated from each culture as starting material. For RNA, mRNAs were purified with the NucleTrap mRNA midi (Macherey Nagel #740656, Düren, Germany) following the manufacturer's protocol. Reverse transcription was then performed with 100% of the mRNA using PrimeScript RT (Clontech #2680A) according to the manufacturer's protocol. cDNAs were concentrated with the DNA Clean & Concentrator kit (Zymoresearch # D4013, Irvine, CA) and used as templates for PCR. Two rounds of PCR were performed for both DNA and RNA sequencing libraries. In the first round, 100% of the purified genomic DNA or 90% of the cDNA were used as a template for PCR (Titanium Taq, 94°C for 3min followed by 16 cycles of 94°C for 30s, 55°C for 10s and 72°C for 30s followed by 68°C for 2min) using the primers nested PCR_FW and nested PCR_RV. The first PCR was then used as template for a second PCR (Titanium Taq, 94°C for 3min followed by 16 cycles of 94°C for 30s, 68°C for 20s and 72°C for 30s followed by 68°C for 2min) using an equimolar mix of the 5 Library-i5_FW primers (see primer table) and the Lib-RV Primers. Each sample has different indexes to allow multiplexing with AMPure XP (Agencourt #A63880) according to the manufacturer's protocol. A first purification with a DNA:Beads ratio of 0.6 was used to remove DNA fragments with a size over 450bp and a second purification with a DNA:Beads ratio of 0.2X removed DNA fragments below 250bp. The purity of the sequencing libraries was verified on a 1% agarose gel and 3-10nM of the cleaned library was sent for massively sequencing using an Illumina HiSeq 4000 sequencer for Paired-End 150 as well as Single-End 50.

#### Read Mapping and Alignment

When reads were at least 100bp long, the bbmerge function available in the BBMap package [9] was used to merge the overlapping paired-end reads (with parameters forcetrimleft=0 qtrim=r minlength=90 maxlength=215 mininsert=190 verystrict=t minoverlap=20).

The adapter sequence was cut out from the 5′ end of the merged read sequence using Cutadapt v1.9 [10] (with parameters -g ACTACGCGTGTTTGAG). The trimmed sequencing read was mapped to the sequence reference using STAR v2.5.3a [11] in local alignment mode, without allowing deletions and insertions or multi-mapping (using parameters –clip5pNbases 0 28 –alignEndsType EndToEnd –alignEndsProtrude 50 ConcordantPair –outFilterMultimapNmax 1 –alignIntronMax 1 –alignIntronMin 2 –outFilterMatchNminOverLread 0.8 –outSAMattributes All).

The resulting SAM files were then processed using an in-house python script to count the number of uniquely mapped reads associated with each sequence and each clone. The last 8 bases at the 5′ end of each aligned read sequence was used to extract the clone ID. When read mates were not merged into one sequence before alignment, only read mates mapping to the same sequence were kept.

To set a minimum threshold for the data quality of a 3′ UTR sequence, we required sequences to be represented by at least 5 clones with more than 5 counts of DNA each, and for each sequence to have at least 1 count of RNA in at least 1 clone.

#### Barcode Misreads

Since MPRAudit relies on the variation from clone to clone, we sought to limit the number of clones erroneously created by misread barcodes. We found that pairs of barcodes with Hamming distance 1 were over-represented for barcodes of the same sequence (but not for barcodes of different sequences). To address this problem, we combined the counts of any two barcodes with Hamming distance one into a single barcode, starting with highest count barcode and keeping the barcode with more counts.

To account for GC-content in the fast-UTR data, we fit a 5th-order polynomial to the steady state expression and stability as a function of GC-content, and use the residuals (plus an offset to set the lowest value to zero). This decision was made by assessing the performance of Nth order polynomial fits ($N \in \{1,...,7\}$) through leave-one-out cross validation.

#### Alternative Prediction Methods

The rules for ARE categories and prediction methods are detailed below. Unless stated otherwise, a sequence is classified by its longest ARE if more than one is present:

- ARED: We follow the rules set out in [12] to create training and test categories. Note that in this publication and the more recent one, many shorter motifs (AUUUA for instance) were excluded from categorization.
- ARED-Plus: We follow the rules set out in [4] to create training and test categories.
- Naive Pentamers: We classify a sequence by the longest complete consecutive AUUUA pentamer (count the number of A's present in the ARE and subtract 1). For instance, "UUUAUUUAUUU" and "AUUUA" would be in the same category.

- Naive Pentamers + 1 Mismatch: We classify a sequence by the longest AUUUA pentamer as above, allowing for a one nt mismatch anywhere.

- Naive AUUUA Count: We classify a sequence by the total number of times the sequence "AUUUA" occurs, including overlaps. If multiple AUUUAs are present we count all of them. For instance, "CCAUUUAUUUAGGAUUUA" has 3.

- Naive Effective Length Pentamers: Pentamers classified by the effective length according to the formula floor((length + registration-2)/4) instead of number of complete AUUUA pentamers. No mismatches allowed.

- 17×4 values of {Length,Registration}: We create a separate category from every possible pair of ARE length and registration values.

- Effective Length (nt): AREs classified by the effective length (length + starting registration), rather than class of pentamer. No mismatch allowed. An out-of-chromosome correction for the small additional y-offset between registrations leads to a small improvement, $\leq 1\%$ of the reported correlation.

- Effective Length (Ending Registration): Using ending registration to calculate effective length rather than starting registration.

- 3 Parameter Regression: A 3-parameter linear fit to the heatmaps in Figures 3F and G (or 4 and 5). The first parameter is the slope with respect to length, the second is the slope with respect to registration, and the third is an offset.

- 2 Parameter Regression: A 2-parameter linear fit (slope and y-intercept) to the aggregated effective length.

- Effective Length Allowing 1 Mismatch: AREs classified by the effective length (length + starting registration), allowing one mismatch. The mismatch cannot be the first or last nucleotide in the ARE (so CCCCAUUUAGGGG would be classified as having length 5).

- Lasso K-Mer Regression: Each 160 nt sequence is broken down into a list of 156 5-mers, which is then used to populate a frequency matrix X. X has dimensions (total number of sequences)×(number of possible 5-mers = 1024). The entries of X are the number of times a given 5-mer occurs in a sequence; it may contain columns with no observations. The model we fit is then y = Xb+c. y is the outcome vector with dimensions (total number of sequences)×1, where each entry is the outcome of a given sequence (gene expression or decay time). b is an effect size vector with dimensions (number of possible 5-mers = 1024)×1, where each entry is the effect of a given K-mer on the outcome. From the training data, we use lasso to find b, given X and y, and we apply b to the test data to obtain predictions. Here the training set is not limited to sequences with AREs, and includes every sequence in the full MPRA; but the test set is limited to AREs as defined in Figure 3. To predict the effect of mutations, we simply subtract the predicted expression or decay time of the wild-type from the predicted expression or decay time of the mutant; we do not train and test on the difference data directly. A similar method was implemented in [13].

| ARED | Motif | Number of $3'$ UTR Segments | ARED-Plus | Motif | Number of $3'$ UTR Segments |
|---|---|---|---|---|---|
| Class I | AUUUAUUUAUUUA UUUAUUUA allowing 1 mismatch | 49 | Cluster 5+ | +AUUUAUUUAUUUA UUUAUUUA+ allowing 1 mismatch | 49 |
| Class II | AUUUAUUUAUUUAUUUA allowing 1 mismatch | 93 | Cluster 4 | AUUUAUUUAUUUAUUUA no mismatches allowed | 6 |
| Class III | AUUUAUUUAUUUA allowing 1 mismatch | 674 | Cluster 3 | AUUUAUUUAUUUA no mismatches allowed | 110 |
| Class IV | WWA[UUUAUUU]AWW allowing 1 mismatch outside the bracketed core | 2157 | Cluster 2 | W[UAUUUAUUUA]WW or WW[AUUUAUUUAU]W allowing 2 mismatches outside the bracketed core | 896 |
| Class V | WWWU[AUUUA]UWWW allowing 1 mismatch outside the bracketed core | 6169 | Cluster 1 | WWWW[AUUUA]WWWW ≥2 of positions 3,4,10,11 must be U; only one mismatch allowed at positions 1,2,12,13; and at least one of position 4,10 must be U | 4772 |
| AUUUA* | AUUUA | 1281 | AUUUA* | AUUUA | 4590 |

Table 1: ARE Class and Cluster definitions from ARED and ARED-Plus[14, 4]. * Indicates $3'$ UTR segments containing an AUUUA sequence but not conforming to the class or cluster motifs.

| N | Steady State Corr. | mRNA Stability Corr. |
|---|---|---|
| 1 | 0.3660 | 0.1137 |
| 2 | 0.3820 | 0.1417 |
| 3 | 0.3825 | 0.1705 |
| 4 | 0.3839 | 0.1708 |
| 5 | 0.3840 | 0.1736 |
| 6 | 0.3843 | 0.1736 |
| 7 | 0.3843 | 0.1734 |

Table 2: Correlation between predicted and actual gene expression (through leave-one-out cross validation) for Nth order polynomial predictions of steady state and stability data for Jurkat T cell fast-UTR results. We used N=5 for our analysis since higher order polynomials provide diminishing returns.

| Most Positive 5-Mers | Lasso Amplitude | Number of Observations | Most Negative 5-Mer | Lasso Amplitude | Number of Observations |
|---|---|---|---|---|---|
| GCUUU | 0.0090 | 7883 | CACUA | -0.0194 | 3165 |
| ACCCC | 0.0078 | 2730 | ACACU | -0.0187 | 4478 |
| AUCGC | 0.0076 | 469 | CGCGG | -0.0166 | 227 |
| CCCAC | 0.0074 | 3440 | UGCAC | -0.0161 | 3920 |
| AGCGA | 0.0070 | 715 | AGACU | -0.0156 | 4563 |
| UUAGA | 0.0065 | 6602 | GGACG | -0.0132 | 513 |
| UACGC | 0.0064 | 401 | GAACU | -0.0120 | 4799 |
| GUAGU | 0.0064 | 3598 | **AUUUA** | -0.0120 | 26795 |
| CCAUU | 0.0064 | 5505 | GGUGC | -0.0120 | 2724 |
| CCAAC | 0.0062 | 2480 | GUCGA | -0.0119 | 319 |
| UGGCA | 0.0061 | 4500 | GGACA | -0.0116 | 3214 |
| UAGUC | 0.0060 | 2654 | UGACU | -0.0115 | 5498 |
| CGGCA | 0.0060 | 689 | ACCUC | -0.0115 | 3865 |
| CUUAU | 0.0057 | 6437 | CGGGC | -0.0100 | 679 |
| CAUCA | 0.0055 | 6678 | CGUUC | -0.0099 | 545 |
| UGGUA | 0.0054 | 4106 | GCCGG | -0.0099 | 803 |
| CUCCC | 0.0054 | 5202 | GUACA | -0.0095 | 4180 |
| CGUAU | 0.0052 | 993 | GACGC | -0.0093 | 460 |
| UCCCC | 0.0049 | 4029 | GACAU | -0.0093 | 4038 |
| CCGAA | 0.0047 | 489 | UCGAU | -0.0091 | 520 |
| UUGUC | 0.0047 | 4989 | CGCGU | -0.0086 | 175 |
| CCCGC | 0.0046 | 882 | GCACU | -0.0083 | 3872 |
| CCGCU | 0.0044 | 649 | ACUUU | -0.0079 | 11306 |
| GGUAG | 0.0043 | 2430 | ACUGG | -0.0073 | 3957 |
| GCAGU | 0.0042 | 4689 | ACGUU | -0.0072 | 1315 |
| CUGUC | 0.0041 | 5015 | CGGUU | -0.0070 | 585 |
| UUAGG | 0.0039 | 3964 | GACUC | -0.0069 | 2599 |
| ACAUC | 0.0039 | 3635 | UGGAC | -0.0069 | 3082 |
| CACGC | 0.0038 | 847 | AAACU | -0.0069 | 8985 |
| UGUCC | 0.0038 | 3990 | GGGGC | -0.0064 | 2594 |

Table 3: Lasso amplitudes of the 20 most positive and negative 5-mers for mRNA stability in Jurkat cells

| ARE-like 5-Mer | Lasso Amplitude | Number of Observations | ARE-like 5-Mer | Lasso Amplitude | Number of Observations |
|---|---|---|---|---|---|
| AUUUA | -0.0124 | 26795 | UUACU | 0.0020 | 7679 |
| UUUAU | -0.0047 | 33724 | UUAGU | 0.0004 | 6432 |
| UUAUU | -0.0059 | 229978 | UAUCU | 0.0010 | 14194 |
| UAUUU | -0.0060 | 39915 | UAUGU | -0.0020 | 16635 |
| | | | AUUCA | 0 | 7777 |
| CUAUU | 0 | 8989 | AUUGA | 0 | 6574 |
| GUAUU | 0 | 10421 | UUUCU | 0 | 18378 |
| CAUUU | 0.0025 | 14630 | UUUGU | 0 | 17396 |
| GAUUU | 0 | 8916 | | | |
| CUUUA | 0 | 9437 | UAUUC | 0 | 7172 |
| GUUUA | 0 | 8281 | UAUUG | 0 | 7817 |
| CUUAU | 0.0057 | 6437 | AUUUC | 0 | 10537 |
| GUUAU | 0 | 6332 | AUUUG | 0.0004 | 12133 |
| | | | UUUAC | 0.0017 | 8342 |
| UCAUU | 0.0010 | 10118 | UUUAG | 0.0028 | 8932 |
| UGAUU | 0.0013 | 8218 | UUAUC | 0.0015 | 7780 |
| UCUUU | 0.0008 | 13822 | UUAUG | 0 | 8588 |
| UGUUU | 0 | 19287 | | | |
| ACUUA | 0 | 6003 | | | |
| AGUUA | 0 | 6075 | | | |
| UCUAU | 0 | 15788 | | | |
| UGUAU | -0.0004 | 16246 | | | |
| UUCUU | 0 | 13801 | | | |
| UUGUU | 0 | 13730 | | | |
| UACUU | 0 | 8249 | | Other 5-Mers | |
| UAGUU | 0.0030 | 6627 | | | |
| AUCUA | 0 | 15882 | AAAAA | 0.0005 | 25485 |
| AUGUA | 0 | 16224 | UUUUU | 0.0005 | 46292 |
| UUCAU | 0.0008 | 10860 | CCCCC | 0.0025 | 3097 |
| UUGAU | 0.0030 | 7393 | GGGGG | 0.0008 | 3140 |

Table 4: Lasso amplitudes of ARE, ARE-like, and other sequences in Jurkat cells.

| # | Category or Method | Steady-State Expression | Stability | Δ Steady-State | Δ Stability |
|---|---|---|---|---|---|
| i | ARED | 0.07 | 0.23 | 0.11 | 0.30 |
| ii | ARED-Plus | 0.09 | 0.21 | 0.14 | 0.29 |
| iii | Naive Pentamers | 0.08 | 0.20 | 0.12 | 0.28 |
| iv | Naive Pentamers + 1 Mismatch | 0.07 | 0.24 | 0.10 | 0.28 |
| v | Naive AUUUA Count | 0.06 | 0.18 | 0.06 | 0.17 |
| vi | Naive Effective Length Pentamers | 0.11 | 0.29 | 0.14 | 0.34 |
| vii | 17×4 values of {Length,Registration} | 0.13 | 0.29 | 0.13 | 0.35 |
| viii | Effective Length (nt) (Length + Registration) | 0.14 | 0.30 | 0.14 | 0.36 |
| ix | Effective Length (nt) (Ending Registration) | 0.13 | 0.29 | 0.14 | 0.35 |
| x | 3 Parameter Regression | 0.11 | 0.30 | 0.14 | 0.36 |
| xi | 2 Parameter Regression | 0.13 | 0.29 | 0.14 | 0.34 |
| xii | Effective Length (nt) Allowing 1 Mismatch | 0.08 | 0.29 | 0.08 | 0.24 |
| xiii | Lasso K-Mer Regression | 0.26 | 0.49 | 0.13 | 0.34 |

Table 5: Out-of-chromosome correlations between predictions and measured data for several ARE categories or simple prediction methods. The columns represent (1) gene expression, (2) stability; (3) the change in steady state expression and the change in stability due to deliberate mutations, respectively. Rows give different ARE categorization or prediction methods. The first two methods are existing methods of ARE categorization in the literature; the next three are naive categorization choices. The next several methods make use of both the ARE length and registration as important parameters. The lasso is trained on the full dataset (it is not ARE-specific) but predictions are tested only on sequences with AREs.

| # | Category or Method | Expression at $T_0$ | Decay Time at $T_4/(T_4+T_0)$ | M-R at $T_0$ | M-R at $T_4/(T_4+T_0)$ |
|---|---|---|---|---|---|
| i | ARED | 0.16 | 0.37 | 0.19 | 0.47 |
| ii | ARED-Plus | 0.19 | 0.40 | 0.20 | 0.48 |
| iii | Naive Pentamers | 0.14 | 0.35 | 0.16 | 0.46 |
| iv | Naive Pentamers + 1 Mismatch | 0.17 | 0.34 | 0.18 | 0.47 |
| v | Naive AUUUA Count | 0.11 | 0.21 | 0.11 | 0.26 |
| vi | Naive Effective Length Pentamers | 0.18 | 0.42 | 0.20 | 0.51 |
| vii | 17×4 values of {Length,Registration} | 0.20 | 0.41 | 0.18 | 0.50 |
| viii | Effective Length (nt) (Length + Registration) | 0.22 | 0.43 | 0.21 | 0.52 |
| ix | Effective Length (nt) (Ending Registration) | 0.21 | 0.41 | 0.21 | 0.51 |
| x | 3 Parameter Regression | 0.22 | 0.43 | 0.20 | 0.52 |
| xi | 2 Parameter Regression | 0.21 | 0.41 | 0.20 | 0.49 |
| xii | Effective Length (nt) Allowing 1 Mismatch | 0.08 | 0.29 | 0.08 | 0.24 |
| xiii | Lasso K-Mer Regression Beas2B | 0.22 | 0.52 | 0.12 | 0.36 |
| xiii | Lasso K-Mer Jurkat ⇒ Beas2B | 0.20 | 0.51 | 0.14 | 0.36 |
| xiii | Lasso K-Mer Beas2B ⇒ Jurkat | 0.14 | 0.39 | 0.08 | 0.24 |

Table 6: Out-of-chromosome correlations between predictions and measured data for Beas2B. The columns represent (1) gene expression, (2) stability; (3) the change in steady state expression and the change in stability due to deliberate mutations, respectively. Rows give different ARE categorization or prediction methods. The first two methods are existing methods of ARE categorization in the literature; the next three are naive categorization choices. The next several methods make use of both the ARE length and registration as important parameters. The lasso is trained on the full dataset (it is not ARE-specific) but predictions are tested only on sequences with AREs. In the end we show training and testing on different cell lines: "Jurkat ⇒ Beas2B" means the Lasso was trained on Jurkat data and tested on Beas2B; "Beas2B ⇒ Jurkat" is the reverse.

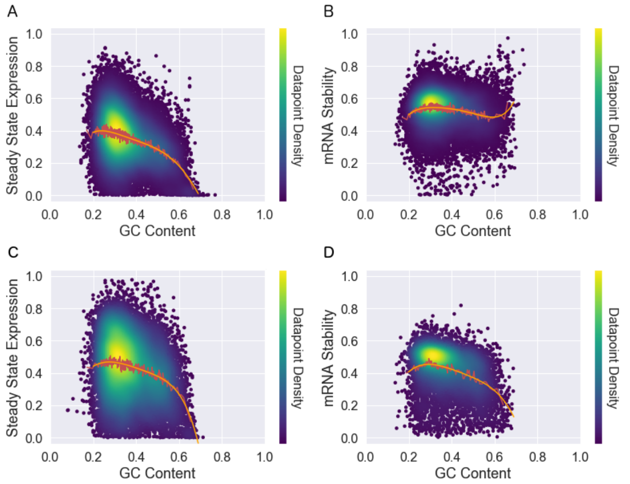

Figure 1: Steady state expression and stability vary as a function of GC content. Each dot represents one sequence, with a yellow-hot color scale given by the density of surrounding data points calculated with a Gaussian kernel and the highest-density points plotted last. The red line is a moving average of 80 3′ UTR segments; the orange line is the 5th-order polynomial fit that we used to residualize the data based on Table 1, which shows minimal improvement with higher-order polynomials. Panels (A-B) show Jurkat data and Panels (C-D) show Beas2B.

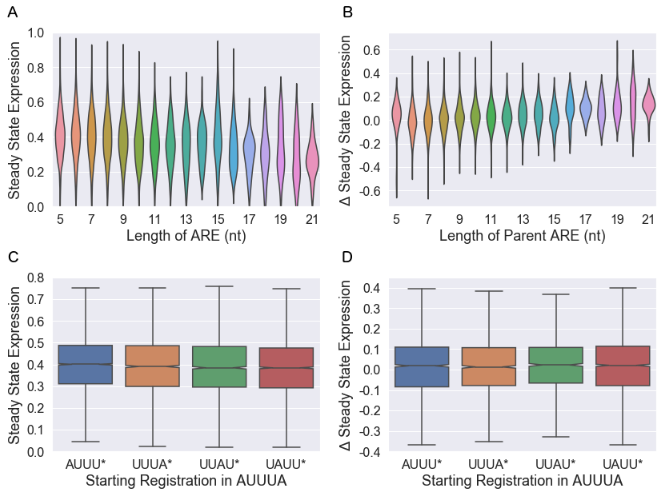

Figure 2: mRNA steady state expression as a function of ARE length and registration for Jurkat data. Linear regression gives two-sided p-values of $4 \times 10^{-67}$, $4 \times 10^{-56}$, 0.23, and $1.5 \times 10^{-8}$, respectively.

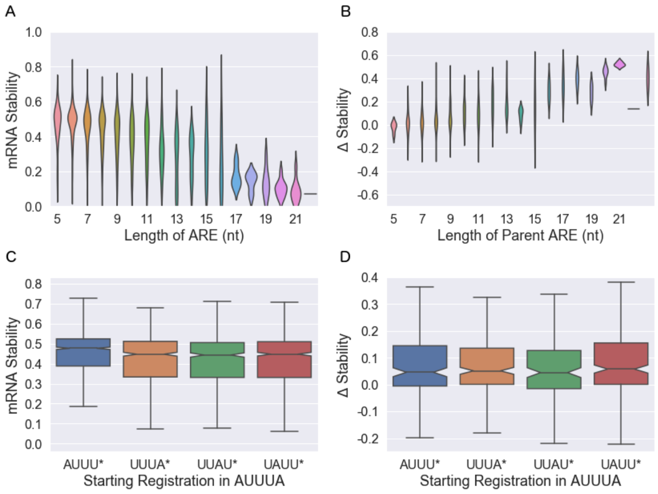

Figure 3: mRNA stability, given as a function of ARE length and registration for Beas2B data. Linear regression gives two-sided p-values of $4 \times 10^{-177}$, $1 \times 10^{-68}$, $4 \times 10^{-8}$, and 0.6, respectively.

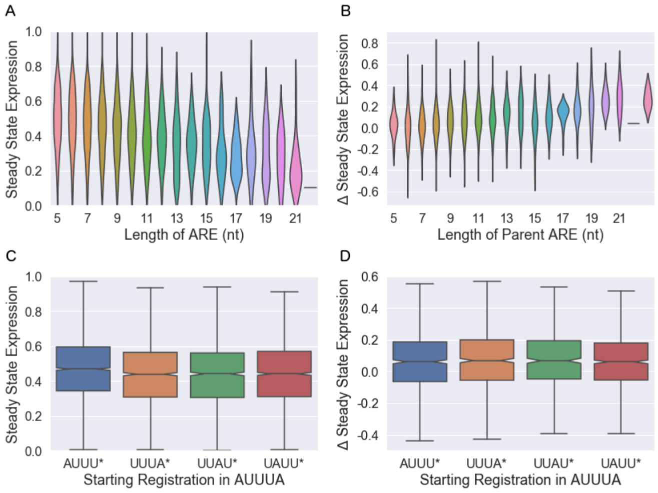

Figure 4: mRNA steady state expression as a function of ARE length and registration for Beas2B data. Linear regression gives two-sided p-values of $8 \times 10^{-96}$, $2 \times 10^{-37}$, $1 \times 10^{-9}$, and 0.9.

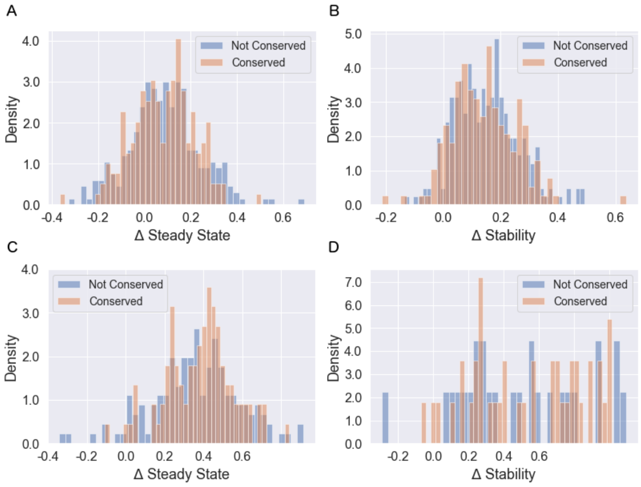

Figure 5: Distribution of conserved and not conserved elements for Jurkat (A-B) and Beas2B (C-D). Panels (A) and (C) show the change in steady state, while Panels (B) and (D) show the change in stability. Comparing these distributions with a Mann-Whitney U test gives p-values of 0.46, 0.24, 0.21, and 0.27, respectively. Within the group of conserved elements, regressing on LOD score gives p-values of 0.36, 0.74, 0.37, and 0.34, respectively.

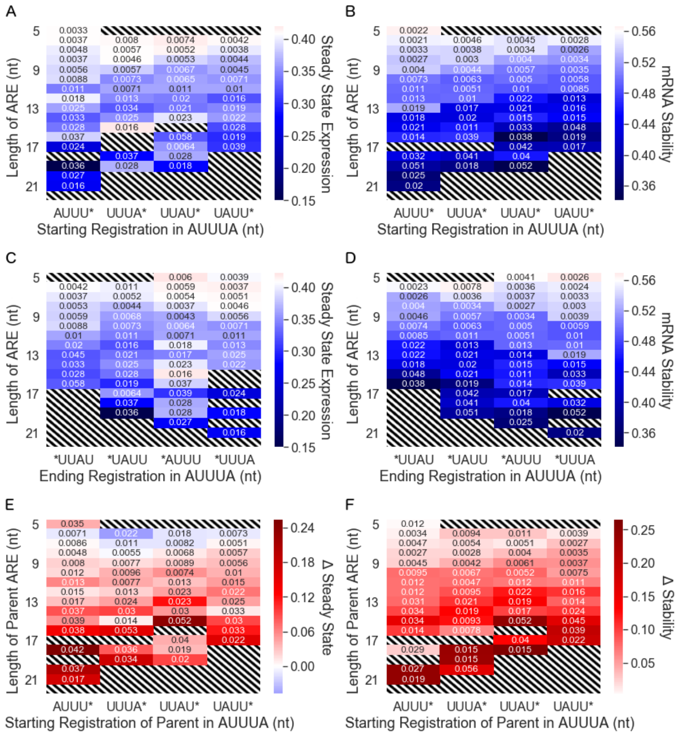

Figure 6: Jurkat data showing mRNA steady state expression and stability, given as a function of ARE length and registration. Panels (A-D) show steady- state expression (A,C) and stability (B,D), respectively, while Panels (E) and (F) show the effect of mutation on steady-state expression and stability, respectively. Values inside of boxes indicate standard errors. The x-axis gives the registration of the start of the ARE in (A,B) and (E,F), and the end of the ARE in (C,D). The y-axis is the length of the ARE, assuming no mismatches. The color scale in (A-D) is determined by setting white to be the mean of all sequences with no ARE, and for (E-F) by setting white to zero.

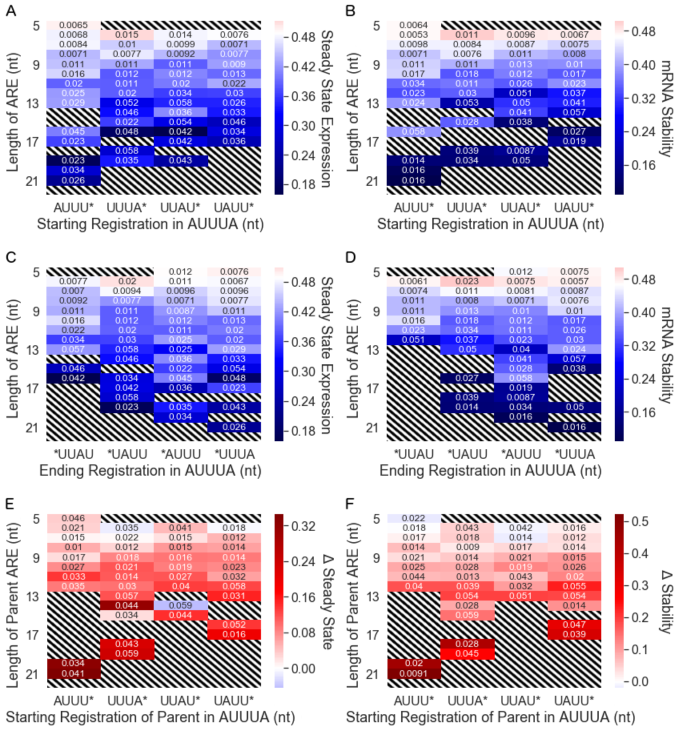

Figure 7: This is the Beas2B version of Figure 6.

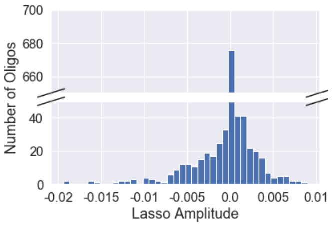

Figure 8: Distribution of Lasso amplitudes for mRNA stability (Jurkat data). The Lasso amplitude is the predicted magnitude of the effect of a 5-mer on mRNA stability; K-mers with positive/negative amplitudes contribute to a relative increase/decrease in mRNA stability. The peak at 0 is a common feature of Lasso regression.

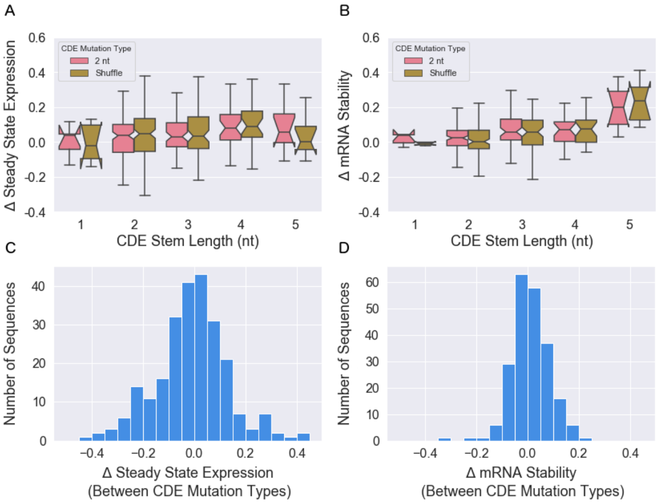

Figure 9: Negligible Difference Between CDE Mutation Types. (A) Change in steady state expression vs CDE stem length, separating 2 nt mutation from full shuffle mutation. (B) Change in stability vs CDE stem length. In (C) and (D), for each reference sequence we subtract the steady state expression and stability (respectively) of the two mutant sequences. (C) The distribution of the difference in steady state expression is centered near zero (0.009±0.010). (D) The distribution of differences in stability between CDE mutation types is also centered near zero (-0.011±0.018). Uncertainty is reported as 95% confidence intervals.

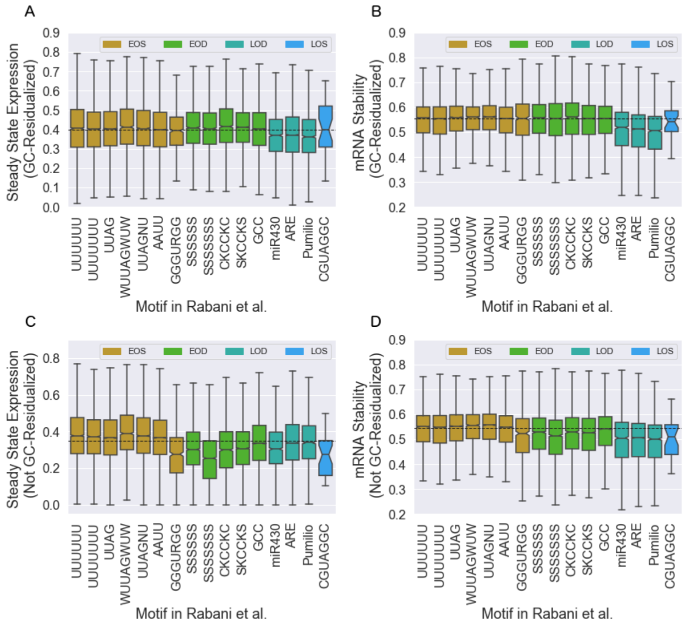

Figure 10: Comparison with Rabani et al. (2017) [13]. Jurkat data summarizing steady state expression (A,C) and mRNA stability (B,D) for the motifs identified in Rabani et al. Boxplots give medians and quantiles, notches denote confidence intervals. The black horizontal dashed line denotes the grand median over all sequences in this study. Colors denote sequence motifs expected to be "early-onset stabilizing" (EOS), "early-onset destabilizing" (EOD), "late-onset destabilizing" (LOD), "late-onset stabilizing" (LOS), respectively, as defined in the reference. Panels (A,B) are GC-residualized to remove the effects of GC-bias, while panels (C,D) are not. Most of these sequences have very similar effects as the dataset median on average once GC-bias is accounted for, although miR430, ARE, and Pumilio sequences are notable exceptions.

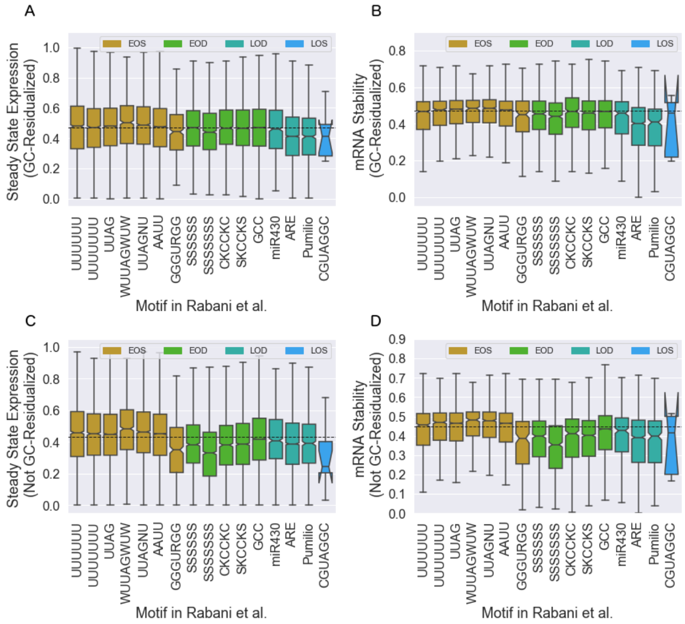

Figure 11: Comparison with Rabani et al. (2017) [13], Beas2B data version of Figure 10.

### References

[1] Zhao, W. et al. Massively parallel functional annotation of 3′ untranslated regions. Nature Biotechnology **32**, 387–391 (2014).

[2] Consortium, T. . G. P. A map of human genome variation from population-scale sequencing. Nature **467**, 1061–1073 (2010).

[3] Mathias, R. A. et al. A continuum of admixture in the western hemisphere revealed by the african diaspora genome. Nature Communications **7** (2016).

[4] Bakheet, T., Hitti, E. & Khabar, K. S. A. ARED-plus: an updated and expanded database of AU-rich element-containing mRNAs and pre-mRNAs. Nucleic Acids Research **46**, D218–D220 (2017).

[5] Alexeyev, M. F., Fayzulin, R., Shokolenko, I. N. & Pastukh, V. A retro-lentiviral system for doxycycline-inducible gene expression and gene knockdown in cells with limited proliferative capacity. Molecular Biology Reports **37**, 1987–1991 (2009).

[6] Dull, T. et al. A third-generation lentivirus vector with a conditional packaging system. Journal of virology **72**, 8463–8471 (1998).

[7] Barde, I., Salmon, P. & Trono, D. Production and titration of lentiviral vectors. Curr. Protoc. Neurosci. **53**, 4.21.1–4.21.23 (2010).

[8] Zeng, H. et al. Lack of ABCG2 expression and side population properties in human pluripotent stem cells. Stem Cells **27**, 2435–2445 (2009).

[9] Bushnell, B., Rood, J. & Singer, E. BBMerge – accurate paired shotgun read merging via overlap. PLOS ONE **12**, e0185056 (2017).

[10] Martin, M. Cutadapt removes adapter sequences from high-throughput sequencing reads. EMBnet.journal **17**, 10 (2011).

[11] Dobin, A. et al. STAR: ultrafast universal RNA-seq aligner. Bioinformatics **29**, 15–21 (2012).

[12] Bakheet, T. ARED 2.0: an update of AU-rich element mRNA database. Nucleic Acids Research **31**, 421–423 (2003).

[13] Rabani, M., Pieper, L., Chew, G.-L. & Schier, A. F. A massively parallel reporter assay of 3′ UTR sequences identifies in vivo rules for mRNA degradation. Molecular Cell **68**, 1083–1094.e5 (2017).

[14] Bakheet, T. ARED: human AU-rich element-containing mRNA database reveals an unexpectedly diverse functional repertoire of encoded proteins. Nucleic Acids Research **29**, 246–254 (2001).
